# Blockage of Lamin-A/C loss diminishes the pro-inflammatory macrophage response

**DOI:** 10.1101/2022.02.22.481454

**Authors:** Johanna L. Mehl, Ashley Earle, Jan Lammerding, Musa Mhlanga, Viola Vogel, Nikhil Jain

## Abstract

Mutations and defects in nuclear lamins can cause major pathologies in affected tissues. Recent studies have also established potential links between lamins, inflammation, and inflammatory diseases but the underlying molecular mechanisms are unknown. We now report that pro-inflammatory activation of macrophages reduces levels of Lamin-A/C to augment pro-inflammatory gene expression and cytokine secretion. We show that activation of bone-marrow derived macrophages (BMDMs) degrades Lamin-A/C, as preceded by its phosphorylation, which is mediated by Caspase-6 and CDK1, respectively. Inhibiting Lamin-A/C phosphorylation and degradation severely inhibits pro-inflammatory gene expression and cytokine secretion. Using LPS-activated Lamin-A/C Knock Out BMDMs, we confirmed that the activation of the IFN-β-STAT pathway is amplified due to Lamin-A/C reduction, which ultimately augments the pro-inflammatory response. As Lamin-A/C is a previously unappreciated regulator of the pro-inflammatory macrophage response, our findings could provide novel opportunities to treat inflammatory diseases. In first proof-of-concept studies we show that macrophage pro-inflammation, as induced by Lipopolysaccharide or pathogenic E. coli, could be reduced by inhibiting Lamin-A/C phosphorylation and degradation. The inhibition of macrophage pro-inflammation could also be achieved by inhibiting members of the Lamin-A/C regulated IFN-β-STAT pathway, i.e., phospho-STAT1 and phospho-STAT3. This newly found mechanism to suppress the pro-inflammatory response of macrophages will provoke a re-thinking of how inflammation can be addressed therapeutically.

## Introduction

While the cell nucleus serves as the site of signal processing and transcription, the nuclear envelope, comprising the nuclear lamina, along with other cellular components(1) regulates the signal transmission from the cytoplasm to the nucleus and the structural organization of the chromatin(2). The nuclear lamina is composed of a dense meshwork assembled from nuclear A-type lamins (Lamin-A and C), B-type lamins (Lamin-B1 and-B2) and associated proteins and located on the inner side of the nuclear envelope(3, 4). Along with providing mechanical stability to the nucleus, nuclear lamina also plays a central role in the 3-dimensional organization of chromatin, and thereby regulates gene expression profiles by modulating the activity of chromatin remodeling enzymes, and transcription factors(5, 6). Owing to its central role in maintaining nuclear architecture and function, it has been shown that perturbations to the nuclear lamina, mutations or reductions, contributes to the broad spectrum of disease manifestations in humans, collectively termed laminopathies(7). Thus, understanding the assembly and disassembly of the nuclear lamina and associated proteins, and underlying molecular mechanisms has become an important biomedical challenge.

The involvement of nuclear lamina and associated changes in nuclear architecture in the immune response was implicated recently(8). For instance, T-cell activation results in a significant upregulation of Lamin-A, which is required for activation of the T cell receptor(9, 10). Mutations in *Lamin-A* gene have been linked to an abnormal increase in inflammation in progeria and laminopathy patients(11). Although it has been suggested that nuclear lamins may be an important mediator in the coordination of immune responses(12–14), as the serum levels of inflammatory cytokines like IL-6, and TNF-α are increased in *Lamin-A* mutant mice(14), a regulatory relationship between nuclear lamins, and an immune cell inflammatory state remains unexplored. Finding novel molecular mechanisms that control inflammatory processes to reduce chronic inflammation, and to improve resolution of inflammation are destined to increase the variety of therapeutic strategies and molecular targets.

When asking whether the nuclear lamins might play a crucial role during the inflammatory activation of macrophages, we discovered that macrophage pro-inflammatory activation results in a significant reduction of Lamin-A/C mRNA and protein levels. Towards gaining mechanistic insights, we found that macrophage pro-inflammatory activation uses machinery involved in mitosis(15–17) to phosphorylate Lamin-A/C protein by CDK1 and to degrade Lamin-A/C protein by Caspase-6, leading to an overall reduction of Lamin-A/C levels. During mitosis, nuclear lamina gets dissolved due to phosphorylation of Lamin protein by CDK1, as phosphorylated Lamin cannot polymerize to form nuclear lamina meshwork(18). However, unlike during mitosis, Lamin-A/C phosphorylation does not lead to a complete breakdown of the nuclear envelope during macrophage pro-inflammatory activation. This decrease in Lamin-A/C levels during macrophage activation is opposite to the Lamin-A/C increase observed during T-cells activation(9, 10), suggesting a potential and unexplored opposing role of Lamin-A/C levels during macrophage and T-cell inflammatory activation, which needs further clarification. Functionally, Lamin-A/C reduction also augmented pro-inflammatory gene expression and cytokine secretion.

Revealing whether nuclear lamins might play regulatory roles in pro-inflammatory macrophage activation, and perhaps also in the resolution of inflammation, is not only essential to advance our basic science knowledge but might also provide opportunities to develop novel therapeutic strategies to treat macrophage induced inflammatory diseases. Using specific pharmacological inhibitors against kinases and enzymes responsible for Lamin-A/C phosphorylation and degradation respectively, which are under clinical trials, we asked whether Lamin-A/C reduction is necessary for inflammatory response

Finally and towards mechanistic understanding, despite our expanding knowledge of macrophage pro-inflammatory activation and involved molecular players(19–22), clear limitations remain as macrophage inflammatory onset and progression are tightly co-regulated by many diverse transcriptional and epigenetic processes. We found that Lamin-A/C reduction impinges on enhanced gene expression and secretion of pro-inflammatory cytokines via the augmentation of the IFN-β-STAT pathway. Lastly, as a proof-of-concept, we explored whether bacterial infection also follows the same route of Lamin-A/C reduction for pro-inflammatory macrophage activation or not. Validating the clinical relevance of our findings, we show that the pharmaceutical inhibition of Lamin-A/C reduction can tame the inflammatory response of macrophages when exposed to heat-killed and active *E. coli*.

Our findings show a completely unexpected behavior whereby macrophages, mitotic or non-mitotic, use the same mitotic and apoptotic machinery, but to promote inflammation. It also underscores the importance of Lamin-A/C for macrophage pro-inflammatory activation and identifies Lamin-A/C as a previously unappreciated regulator of the immune response. Our findings suggest a fundamentally new understanding of a so far little explored mechanism by which the nuclear lamina regulates macrophage pro-inflammatory response. This may enable the future development of drugs for treating diseases associated with excessive secretion of pro-inflammatory cytokines and of other macrophage-derived metabolites, such as chronic inflammatory diseases, by targeting Lamin-A/C phosphorylation, degradation and associated molecular mechanisms.

## Results

### Macrophage pro-inflammatory activation results in Lamin-A/C downregulation

To examine whether macrophage pro-inflammatory activation alters Lamin-A/C expression *in vivo*, we first re-examined a previously published RNA-Sequencing data set of microglia from Lipopolysaccharide (LPS) injected wildtype (WT) mice, where microglia were isolated 24 hours post LPS-injection (Fig. 1A)(23). As expected, the pro-inflammatory genes were highly upregulated, including those encoding for cytokines (e.g., *Il6, Il10, I12b* and *Tnf-α*), chemokines (e.g., *Ccl4, Ccl3* and *Ccl5*) and enzymes necessary for pro-inflammatory marker synthesis (*Nos2* and *Ptgs2*). Yet, the same data set also revealed that the mRNA expression of *Lamin-A/C* in these pro-inflammatory activated microglia was significantly downregulated (Fig. 1A). When re-examining other previously published RNA-Sequencing datasets of *in vitro* LPS activated tissue resident macrophages, we found that the same correlation holds true for macrophages from different organs in mice and humans-peritoneal(24), microglia(25), bone marrow derived macrophages (BMDMs) from two different mouse strains(26), as well as for human monocytes(27) and human alveolar macrophages(28) (Fig. 1A). The same holds true for LPS-activated BMDMs across species, including rat, buffalo, cow, goat and horse(29) (Figs. 1A&S1) and for two different macrophage cell lines J774A.1 (mouse)(30) and THP-1 (human)(31) (Fig. 1A). In all these described data sets, macrophages were isolated from the respective tissues, cultured according to the respective protocols, and treated with LPS for 2 to 6 hours(24–31). Remarkably, the downregulation of *Lamin-A/C* mRNA levels occurred universally in all species, cell lines, and primary tissue resident macrophages, without any sexual dimorphism, *in vivo* and *in vitro* (Figs. 1A&S1). This suggests a previously unknown inverse-correlation between pro-inflammatory genes and *Lamin-A/C* mRNA expression levels. It should be noted that all of this was done on mRNA level, and that effect on Lamin-A/C protein level needs to be determined.

**Fig. 1:**
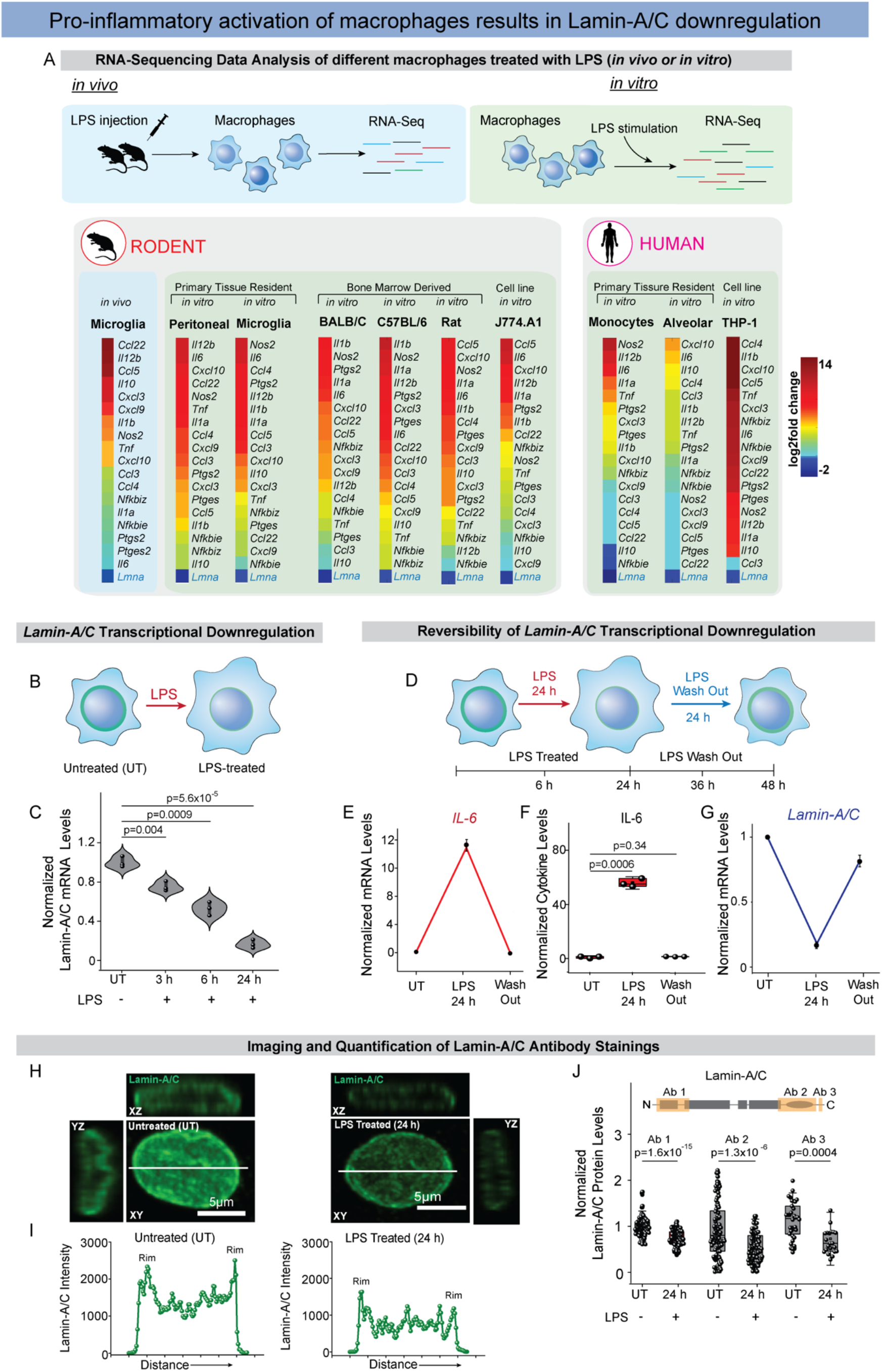
Pro-inflammatory macrophage activation results in a *Lamin-A/C* downregulation in mice and human, independent of the organ or gender of the donors: **(A)** Color coded map shows the mRNA expression levels of various pro-inflammatory genes and *Lamin-A/C* in microglia of LPS-injected mice(23), in LPS-treated mouse peritoneal macrophages(24), microglia(25), bone marrow derived macrophages isolated from rodents of different species (mouse and rat(29)) and mouse strains(26) (BALB/C and C57BL/6), as well as in LPS treated human monocyte(27) and alveolar macrophage(28). The same is seen in LPS treated macrophage cell line J774.A1 (mouse)(30) and THP-1 (human)(31). Expression data were obtained from public repositories as cited. **(B)** Scheme of experimental setup, BMDMs were treated with LPS for different durations of time. **(C)** Violin plots show the normalized levels of *Lamin-A/C* mRNA in Untreated (UT), and LPS treated BMDMs as determined by qPCR. Levels were further normalized to UT BMDMs conditions to find the fold change. **(D)** Scheme of experimental setup to illustrate the reversibility of the process. BMDMs were treated with LPS for 24 h for pro-inflammatory activation followed by culturing in media without LPS for 24h (Wash out) to shed light on inflammation and resolution in macrophages. **(E)** Time course quantification shows a reversal in the gene expression levels of pro-inflammatory gene *IL-6* in UT, 24 h LPS treated BMDMs and after 24 h LPS Wash out. Data are presented as Mean±S.E and normalized to UT BMDMs condition to find the fold change. **(F)** Box plots show reversal of secreted IL-6 levels in UT, 24h LPS treated BMDMs and after 24 h LPS Wash out. Levels were normalized to UT BMDMs conditions. **(G)** Time course quantification shows a reversal in the expression levels of *Lamin-A/C* in UT, 24h LPS treated BMDMs and after 24h LPS Wash out. Data are presented as Mean±S.E and normalized to UT BMDMs condition to find the fold change. **(H)** Representative orthogonal views of nucleus in UT and LPS treated (24 h) BMDM nuclei stained with Lamin-A/C (green) antibody. Scale Bar = 5 μm. **(I)** Line-intensity profiles (corresponding to white lines in H) of Lamin-A/C in UT, and 24h LPS treated BMDMs. **(J)** Box plots show normalized protein levels of Lamin-A/C in UT, and LPS treated (24 h) BMDMs determined using antibodies binding to different epitopes of the Lamin-A/C protein, as shown in the cartoon. Levels were normalized to UT BMDMs condition. For all the plots *p*-values were obtained with the two-sided Student’s *t*-test. In all the box plots, the boxes show 25th and 75th percentiles, the middle horizontal line shows the median, small open squares show the mean, and whiskers indicate S.D. All the experiments were independently repeated three or more times.

### Downregulation of *Lamin-A/C* mRNA expression levels correlates inversely with the expression of pro-inflammatory genes

To confirm the observed decline in the *Lamin A/C* mRNA expression levels, we stimulated BMDMs with LPS and performed quantitative PCR (qPCR). As early as 3 h after LPS treatment, levels of *Lamin-A/C* mRNA declined by 40% and after 24 h a steady decline to as much as 80% of gene expression was observed (Figs. 1B&C). We further confirmed this using LPS treated alveolar macrophages, which also show a significant reduction in *Lamin-A/C* mRNA levels which inversely correlated with an upregulated pro-inflammatory gene expression (Fig. S2A). Only one exception to this observed inverse correlative relationship between *Lamin-A/C* downregulation and the pro-inflammatory gene expression of macrophages could be identified here, i.e. for the immortalized RAW264.7 macrophage cell line when examining previously published RNA-Sequencing studies(32, 33). Indeed, we were able to confirm that LPS activation of RAW264.7 macrophage results in an increase in Lamin-A/C mRNA level (Fig. S2B), which conflict with our observations made in different primary tissue resident macrophages and in other immortalized macrophage cell lines (Figs. 1A, C&S1). However, this contradictory observation might not be surprising as multiple papers highlight significant cell signalling differences between primary and RAW264.7 macrophages(34, 35). Altogether, our data (Fig. 1C) taken together with the available literature on primary macrophages (Fig. 1A) thus suggest that the previously proposed role of Lamin-A/C during pro-inflammation observed in RAW264.7 macrophages(33) cannot entirely be translated to primary BMDMs and consequently, as Fig. 1A illustrates, to organ-specific tissue resident macrophages *in vivo* and *in vitro*.

Despite the fundamental physiological role of the pro-inflammatory macrophage response to stage a defence against infection, the proper resolution of inflammation is essential for functional tissue repair after pathogen or injury challenge(36). Thus one of the central causes for chronic inflammatory diseases has been assigned to macrophages remaining in a continuous state of pro-inflammatory phenotype(37). To test whether *Lamin-A/C* levels revert to their homeostatic levels during inflammation resolution, we mimicked the resolution phase by first treating BMDMs with LPS for 24 h followed by washing out LPS and culturing BMDMs in media without LPS for another 24 h (Fig. 1D). As expected, there was a sharp decline in transcription of pro-inflammatory genes such as *IL-6* (Fig. 1E) and IL-6 cytokine secretion (Fig. 1F) whereas the mRNA expression levels of *Lamin-A/C* returned to near homeostatic levels (Fig. 1G) post 24 h of LPS washout. These observations further confirm the inverse correlative relationship between *Lamin-A/C* and the pro-inflammatory gene expression.

To check whether Lamin-A/C reduction upon LPS treatment holds true at the Lamin-A/C protein levels as well, BMDMs were treated with LPS (for 24 h) and stained for Lamin-A/C (Fig. 1H). Quantitative fluorescence imaging demonstrated a significant decline in Lamin-A/C protein levels, as detected by antibody staining (Figs. 1I &S3), of more than 50% upon 24 h LPS treatment (Fig. 1J). To exclude the possibility that the decline of the Lamin-A/C levels is owed to the inaccessibility of epitopes due to structural changes in the lamina meshwork(38), and to validate that the observed reduction of antibody binding can be attributed to an authentic Lamin-A/C protein decline, we used multiple antibodies to stain for Lamin-A/C by targeting either its N or C termini (Fig. 1J) and observed a similar and significant decline in Lamin-A/C levels.

### Lamin-A/C protein downregulation in macrophages is accompanied by its degradation

To distinguish between the potential effects on Lamin-A versus Lamin-C, a time course study using immunoblotting was performed that showed a significant reduction in the levels of both isoforms within 6 h of LPS treatment, which were further reduced to almost ∼50% within a 24 h LPS treatment (Figs. 2A&B). To further strengthen the hypothesis that a reduction in Lamin-A and C levels is a general hallmark of pro-inflammatory activation, we treated BMDMs with TNF-α (another pro-inflammatory phenotype inducing chemical agent) (Fig. S4A) and observed a similar decrease of Lamin-A and C protein levels.

**Fig. 2:**
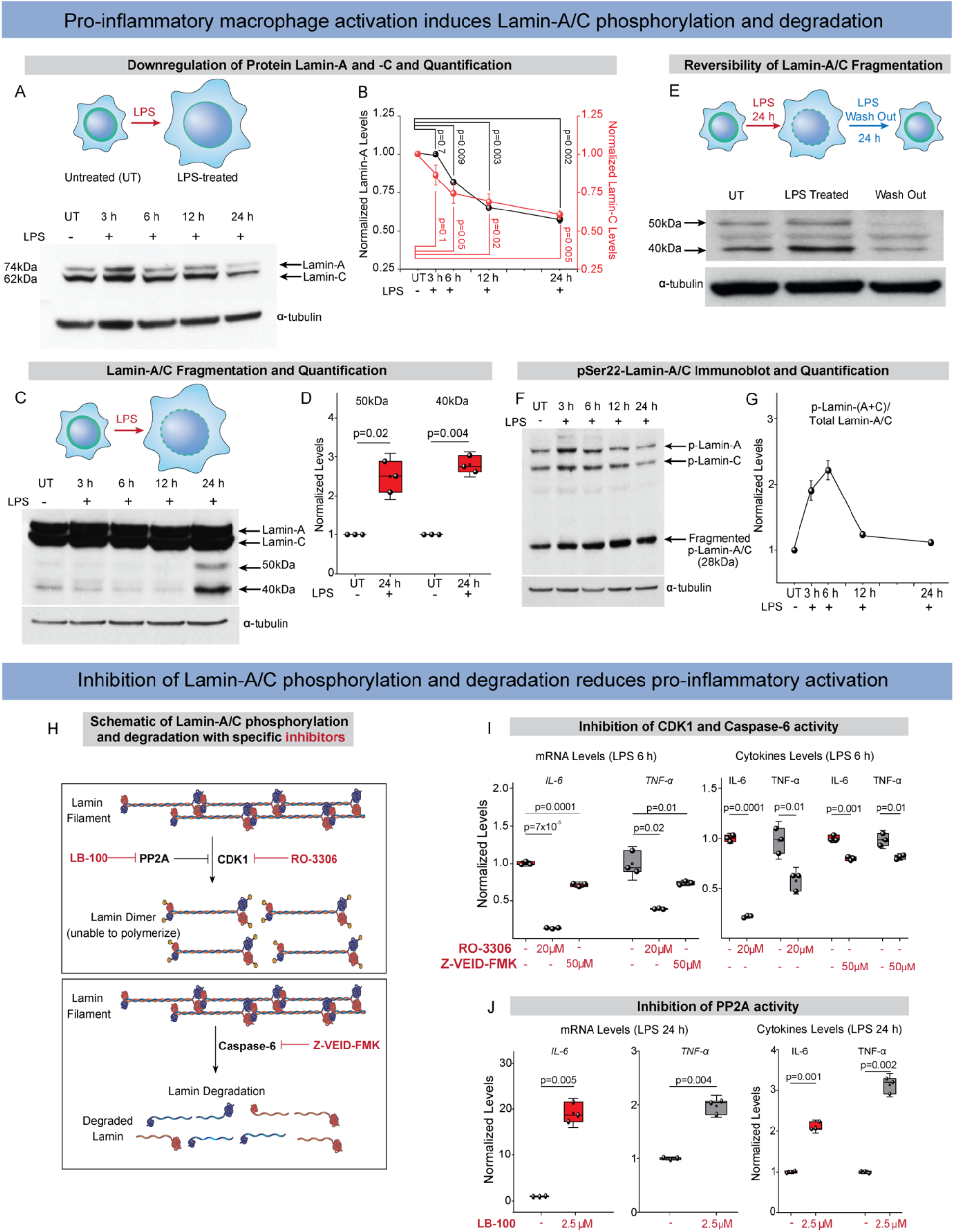
Pro-inflammatory macrophage activation induces Lamin-A/C phosphorylation and degradation: **(A)** Immunoblot shows total levels of Lamin-A and Lamin-C in Untreated (UT), and LPS treated BMDMs. α-tubulin served as a loading control. **(B)** Time course quantifications of Lamin-A and Lamin-C normalized with α-tubulin over three independent immunoblotting experiments. Calculated values were further normalized to the UT BMDMs condition. Data are presented as Mean±S.E. **(C)** Longer exposure of immunoblots against Lamin-A/C in UT, and LPS treated BMDMs to detect lower fragmented Lamin-A/C bands (50kDa and 40kDa). α-tubulin served as a loading control. **(D)** Box plots show degraded Lamin-A/C fragments (50kDa and 40kDa) normalized to α-tubulin between UT and LPS treated BMDMs. Calculated values were further normalized to the UT BMDMs condition. **(E)** Immunoblots against Lamin-A/C in UT, LPS treated, LPS wash-out BMDMs to detect fragmented Lamin-A/C bands (∼50kDa and ∼40kDa). α-tubulin served as a loading control. **(F)** Immunoblot shows total levels of p-Ser22-Lamin-A and p-Ser22-Lamin-C along with a phosphorylated fragmented band of Lamin-A/C (∼ 28kDa) in UT, and LPS treated BMDMs. α-tubulin served as a loading control. **(G)** Time course quantification of total p-Ser22-Lamin-A+C normalized to the total Lamin-A/C over three independent immunoblotting experiments. Calculated values were further normalized to the UT BMDMs condition. **(H) (Top)** Schematic shows the CDK1 mediated Lamin-A/C phosphorylation, which is known from other cells to be blocked by PP2A during cell cycle. Also shown are the drugs used in this study to specifically inhibit the activity of the respective enzymes. **(Bottom)** Schematic shows Caspase-6 mediated Lamin-A/C degradation as inhibited by Z-VEID-FMK. **(I)** Box plots show the lowering of pro-inflammatory gene expression (qPCR) **(Left)** and secreted cytokine levels (ELISA) **(Right)** in LPS treated BMDMs, also treated with RO-3306 or Z-VEID-FMK to inhibit Lamin-A/C phosphorylation and degradation, respectively. Levels were normalized to 6 h LPS treated BMDMs condition to find the fold change. **(J)** Box plots show an increase in the pro-inflammatory gene expression (qPCR) **(Left)** and secreted cytokine levels (ELISA) **(Right)** in LPS treated BMDMs, also treated with LB-100, a PP2A inhibitor, as compared to BMDMs treated with only LPS for 24 h. Levels were normalized to 24 h LPS treated BMDMs conditions to find the fold change. For all the plots *p-*values were obtained with the two-sided Student’s *t*-test. In all the box plots, the boxes show 25th and 75th percentiles, the middle horizontal line shows the median, small open squares show the mean, and whiskers indicate S.D. All the experiments were independently repeated three or more times.

Previous studies have quantified the turnover rates of lamins in quiescent fibroblasts, uncovering that around 10% of Lamin-A/C is replaced in the lamina meshwork with newly synthesized Lamin-A/C proteins within 24 h(39). Since these Lamin-A/C turnover rates are not sufficient to explain the decrease of Lamin-A/C protein levels of up to 50% in LPS treated BMDMs, an active process to more rapidly reduce or downregulate Lamin-A/C protein levels must take place. Interestingly, higher exposed immunoblots showed a significant fragmentation of Lamin-A/C (∼50 kDa and ∼40 kDa) in 24 h LPS treated macrophages (Figs. 2C&D), suggesting their degradation. This also suggests a potential combinatorial effect of both in reducing the total A/C levels during the LPS-treatment (Figs. 1H-J,2A&B), i.e., lower transcription (Fig. 1C) and enhanced degradation of Lamin-A/C (Fig. 2C&D). Interestingly, along with the reversal in the expression of pro-inflammatory genes and *Lamin-A/C* mRNA levels during inflammatory resolution (Figs. 1D-G), 24 h LPS washout reduces the levels of degraded Lamin-A/C fragments (Fig. 2E). These observations suggest a restoration of the Lamin-A/C levels upon LPS withdrawal, strengthening our hypothesis of a potential role of Lamin-A/C during pro-inflammatory activation of macrophages.

### Lamin-A/C degradation is preceded by Lamin-A/C phosphorylation

The compelling observation that Lamin-A/C protein levels are reduced, and Lamin-A/C protein is degraded during LPS treatment motivated us to explore in more depth the events that lead to Lamin-A/C fragmentation. As Lamin-A/C phosphorylation by CDK1 (at residues Serine 22 and 392) triggers Lamin-A/C depolymerization at the onset of nuclear envelope breakdown during mitosis in other cell types(40), and precedes the enzymatic cleavage of Lamin-A/C(41)mostly due to Caspase-6 enzyme activity(42), we conducted time course immunoblotting experiments with BMDMs of phospho-Serine 22 Lamin-A/C (pSer22-Lamin-A/C). Quantification of immunoblots show a significant increase in the pSer22-Lamin-A/C/total Lamin-A/C ratio, peaking at 6 h of LPS-treatment, and then declined as the total pSer22-Lamin-A/C levels diminished over the subsequent 12 and 24 h timepoints (Figs. 2F&G). Confocal microscopy imaging further revealed significant nucleoplasmic accumulation of pSer22-Lamin-A/C upon 6 h LPS treatment (Figs. S4B&C), followed by a decline at 24 h of LPS-treatment (Fig. S4B). We also examined the levels of active CDK1 (phospho-Threonine 161 CDK1) in BMDMs during the first 6 h of LPS treatment by immunoblotting (Fig. S4D). The phosphorylated CDK1 fraction increased significantly within 1 h post-LPS treatment, peaked at 2 h and then returned to homeostatic levels by the end of 3-6 h (Figs. S4D&E), confirming significant activation of CDK1 post LPS treatment which in turn induces Lamin-A/C phosphorylation.

### Pharmacological inhibition of Lamin-A/C phosphorylation and degradation reduces expression of pro-inflammatory genes and cytokine secretion

To verify whether Lamin-A/C phosphorylation and degradation by the enzymes CDK1 and Caspase-6, respectively, is required for pro-inflammatory gene expression, these two enzymes were targeted by RO-3306 (a specific ATP-competitive inhibitor of CDK1)(43) and Z-VEID-FMK (a specific irreversible inhibitor of Caspase-6)(42), respectively (Fig. 2H). RO-3306 is one of many Cyclin-dependent kinase inhibitors, which have been identified as potential anti-inflammatory agents(44) and in human phase-II clinical trials to treat adjunctive therapy for rheumatoid arthritis(45). Z-VEID-FMK is one of multiple caspase inhibitors which have been tested in various preclinical studies for the treatment of cell death related pathologies(46).

Immunoblotting of LPS and RO-3306 or Z-VEID-FMK treated BMDMs indeed confirmed the efficacy of RO-3306 and Z-VEID-FMK in inhibiting either Lamin-A/C phosphorylation (Fig. S5A) or degradation (Fig. S5B), respectively. Strikingly, inhibiting either Lamin-A/C phosphorylation or degradation significantly decreased *IL-6* and *TNF-α* mRNA expression, as well as IL-6 and TNF-α cytokine secretion (Fig. 2I). This shows for the first time, to the best of our knowledge, that Lamin-A/C reduction is required for sufficient pro-inflammatory gene expression and thus for an efficient pro-inflammatory response of BMDMs.

Conversely, to keep Lamin-A/C phosphorylated which can be accomplished by keeping CDK1 constitutively active, we restrained protein phosphatase 2A (PP2A) activity (Fig. 2H)(47), using LB-100, a PP2A specific competitive inhibitor(48) known to have anti-cancer activity(49) (Fig. 2H). PP2A is a mitosis-regulating phosphatase during the cell cycle, which dephosphorylates Cdc25, causing a persistent dephosphorylation (Thr-14 and Tyr-15) and hence activation of Cdk1. Indeed, *IL-6* and *TNF-α* mRNA expression levels were significantly increased, as well as the IL-6 and TNF-α cytokine secretion levels. This crucial experiment further confirms the role of Lamin-A/C phosphorylation and thus reduction in Lamin-A/C levels during BMDMs pro-inflammatory response (Fig. 2J).

### CDK1 and Caspase-6 inhibition has little effect on pro-inflammatory gene expression in Lamin-A/C Knock Out BMDMs

Since pharmacological inhibitors can have unanticipated side effects, we used a mouse model with a deletion in the *Lmna* gene (50) to assess the transcriptional response of pro-inflammatory genes in the absence and presence of LPS, as well as the presence of the inhibitor RO-3306 or Z-VEID-FMK. Absence of Lamin-A/C, post isolation from bone marrow, was confirmed in Lamin-A/C-Knock-Out BMDMs (here after referred as KO-BMDMs) by immunoblotting (Fig. S6A). We also validated that KO-BMDMs differentiate and polarize in the same manner as Wild type (WT) BMDMs, which are known to polarize upon LPS treatment towards M1 phenotype, which is characterized by a significant increase in cell and nuclear spreading areas along with higher cell circularity. The insignificant cellular or nuclear morphological differences between WT- and KO-BMDMs before and after LPS treatment (Figs. S6B-G) confirmed that the KO-BMDMs differentiated normally into functional macrophages. However, KO-BMDMs showed marginally higher cell circularity (Fig. S6F). It is also important to note that Lamin-B1 and Lamin-B2 levels remain unaltered in all the KO-BMDMs(50).

To then test our hypothesis, KO-BMDMs were treated with LPS in the presence or absence of RO-3306. In contrast to WT BMDMs (Fig. 2I), an insignificant downregulation was observed in pro-inflammatory gene expression (*IL-6*) in LPS treated KO-BMDMs (Fig. S5C). Z-VEID-FMK treatment resulted in only a mild upregulation of pro-inflammatory gene expression (Fig. S5C) in LPS treated KO-BMDMs. Taken together, these data show that Lamin-A/C phosphorylation and degradation exerts a previously undescribed role in increasing pro-inflammatory macrophage response, the potential regulatory mechanism of which was probed next.

### Lamin-A/C reduction in Wild Type BMDMs, as maximized in Lamin-A/C-KOs, enhances pro-inflammatory genomic programs mostly belonging to the Interferon-beta pathway

Along with *IL-6* and *TNF-α*, macrophage pro-inflammatory activation leads to upregulation of different pro-inflammatory gene clusters(22). To check if either Lamin-A/C reduction upon LPS treatment, or its elimination has an influence on these other clusters, we performed RNA-Seq experiments using WT and KO BMDMs (Fig. 3A). Use of these KO BMDMs, which asks how pro-inflammatory processes proceed even in the absence of Lamin-A/C, circumvents potential off target effects of inhibitors like RO-3306 and Z-VEID-FMK.

**Fig. 3:**
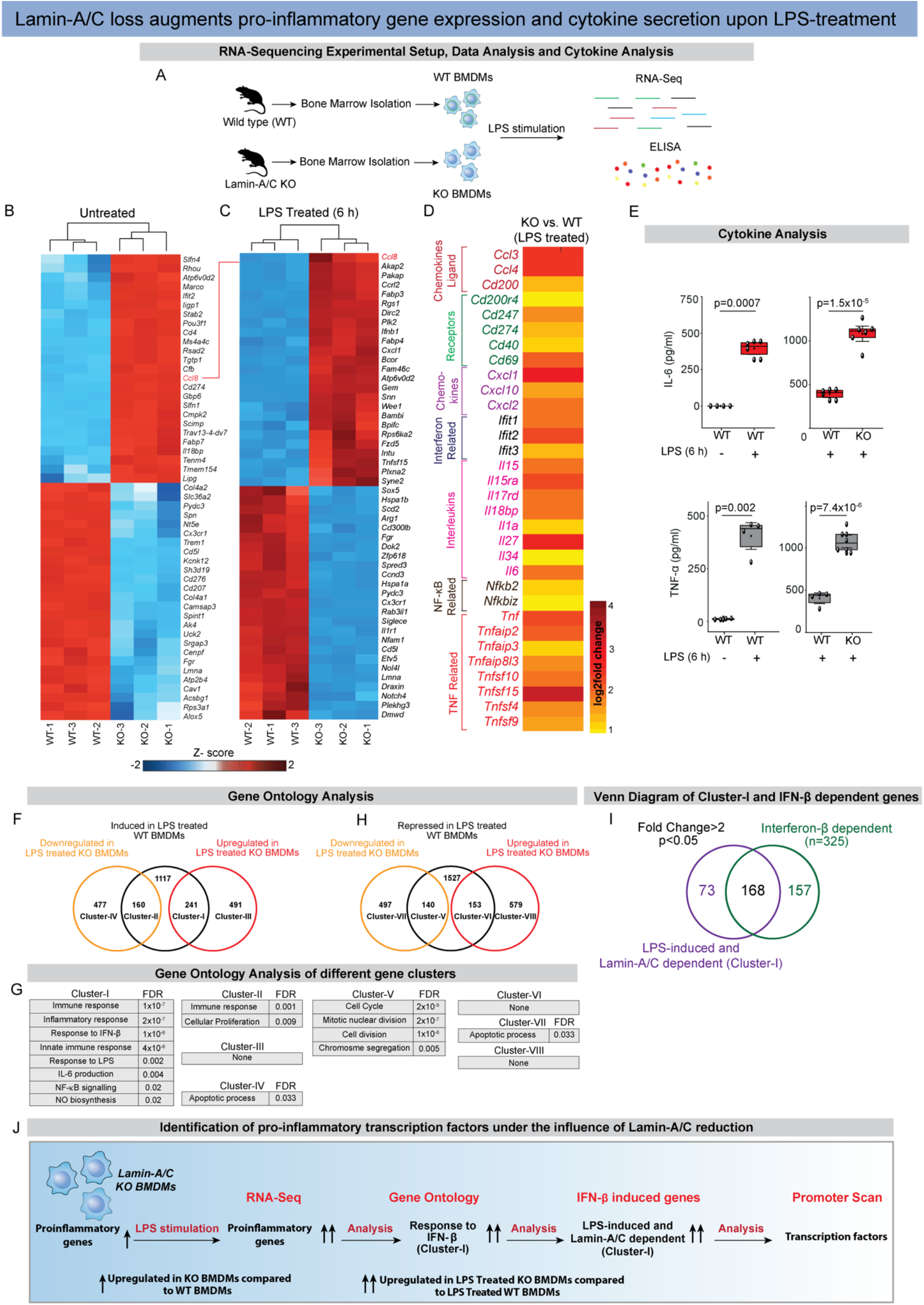
*Lamin-A/C* knockout augments pro-inflammatory genomic programs: **(A)** Scheme shows experimental RNA-Sequencing and ELISA setup between Wild type (WT) and *Lamin-A/C* knockout (KO) BMDMs. **(B)** Heatmap shows the top 25 upregulated and downregulated genes between WT and *Lamin-A/C* KO BMDMs (Fold change>2 and p-value<0.05). **(C)** Heatmap shows the top 25 upregulated and downregulated genes between LPS treated (6 h) WT and KO BMDMs (Fold change>2 and p-value<0.05). (**D)** Heat map shows the levels of selected pro-inflammatory genes upregulated in LPS treated KO-BMDMs as compared to LPS treated WT BMDMs. **(E)** Box plots show the differences in secreted cytokine levels of TNF-α and IL-6 between WT vs. WT+LPS, and WT+LPS vs. KO+LPS treated BMDMs. Levels were normalized to untreated BMDMs condition to find the fold change. **(F)** Venn-diagram shows the number of overlapped genes (Fold change>2 and p < 0.05) between LPS induced genes in WT-BMDMs and genes which get further induced (241) or repressed (160) in LPS treated KO-BMDMs. (**G)** Tables show Gene Ontology analysis of different gene clusters. **(H)** Venn-diagram shows the number of overlapped genes (Fold change>2 and p < 0.05) between LPS repressed genes in WT-BMDMs and genes which further get repressed (140) or induced (153) in LPS treated KO-BMDMs. **(I)** Venn-diagram shows the overlapped genes (Fold change>2 and p < 0.05) between LPS-induced and Lamin-A/C dependent (Cluster-I) and Interferon-β dependent genes. **(J)** Scheme shows experimental setup of RNA-Sequencing and data analysis to identify Lamin-A/C reduction influenced pro-inflammatory transcription factors. For all the plots, *p* values were obtained with the two-sided Student’s *t*-test. In all the box plots, the boxes show 25th and 75th percentiles, the middle horizontal line shows the median, small open squares show the mean, and whiskers indicate S.D. Triplicate samples were used for RNA-Sequencing. All the other experiments were independently repeated two or more times.

Unsupervised cluster analysis segregated WT- and KO-BMDMs and revealed prominent differences between their transcriptomes at baseline (Fig. S7A) and in response to 6 h LPS stimulation (Fig. S7B). Figure 3B shows top 25 up- and down-regulated genes between WT- and KO-BMDMs, suggesting that KO-BMDMs already have higher expression of pro-inflammatory genes like *Ccl8* as compared to WT BMDMs (Fig. 3B). This clearly suggests that pro-inflammatory genomic programs are potentiated in KO-BMDMs. Gene ontology (GO) analysis of WT-vs. KO-BMDMs also suggested that upregulated functional clusters in KO-BMDMs largely belong to the category of inflammation, whereas downregulated functional clusters largely belong to the category of and DNA damage and repair (Fig. S8A).

While looking at differences in LPS treated WT-versus KO-BMDMs, checking the status of *Ccl8* gene suggested that Lamin-A/C reduction also augments LPS response in KO-BMDMs (Fig. 3C). Further analysis of all the genes upregulated in LPS treated KO-BMDMs vs. LPS treated WT-BMDMs revealed several critical and well characterized pro-inflammatory gene clusters (Fig. 3D). These include chemokine related (e.g. *Cxcl1, Cxcl10*), Interferon related (e.g. *Ifit1, 2*, and *3*), Interleukins (e.g. *Il-6, Il-1a*), NF-κB related (e.g. *NF*κ*b2, NF*κ*biz*) and TNF related (e.g. *TNF-α*) genes (Fig. 3D). Not only at the mRNA levels these pro-inflammatory genes were also upregulated at their cytokine secretion levels in LPS treated KO BMDMs (Figs. 3D&E).

To fully characterize these up- and down-regulated genes, and the signaling pathways and biological functions these genes belong to, we further analyzed the RNA-Sequencing data. RNA-Sequencing datasets revealed that 1518 genes were upregulated upon LPS treatment of WT-BMDMs for 6 h, the expression of several of these genes were also dependent on Lamin-A/C reduction (Fig. 3F). 241 genes (Cluster-I) were further upregulated in LPS treated KO-BMDMs, whereas 160 genes (Cluster-II) were downregulated in LPS treated KO-BMDMs (Fig. 3F). GO analysis of Cluster-I gene revealed clusters involved in pro-inflammatory response, innate immune response, response to LPS, response to IFN-β, Nitric Oxide synthesis, and IL-6 productions and several other critical inflammation related clusters (Fig. 3G). Whereas genes in Cluster-II only belonged to immune response and cellular proliferation. GO analysis revealed that genes exclusively upregulated in LPS treated KO-BMDMs (Cluster-III) did not belong to any significant processes, whereas genes exclusively downregulated in LPS treated KO-BMDMs (Cluster-IV) belonged to apoptotic pathways (Fig. 3G). Contrary to this, 1820 genes were downregulated upon LPS treatment of WT-BMDMs for 6 h (Fig. 3H). Several of these genes were further either downregulated (Cluster-V) or upregulated (Cluster-VI) in LPS treated KO-BMDMs (Fig. 3H). GO analysis of Cluster-V revealed processes mainly involved in cell cycle, whereas Clusters-VI did not belong to any major biological processes. GO analysis further revealed that genes in cluster Cluster-VII (exclusively downregulated in KO-LPS treated BMDMs) and Cluster VIII (exclusively upregulated in LPS treated KO BMDMs) did not belong to any major biological processes (Fig. 3G). To understand to which pathways the genes, that were upregulated in LPS treated KO-BMDMs vs. LPS treated WT-BMDMs were associated to, KEGG pathways analysis was performed. Gene sets mainly associated with TNF-α and NF-κB signaling, two central pathways involved in macrophage pro-inflammatory activation (Fig. S8B), were upregulated when comparing LPS treated KO-BMDMs vs. LPS treated WT-BMDMs, whereas the downregulated gene set was mainly associated with the MAP-Kinase signaling pathway (Fig. S8B).

Cluster-I is particularly interesting, as it is the dominant gene cluster involved in regulating the pro-inflammatory response of BMDMs and at the same time is seen to be upregulated by *Lamin-A/C* reduction or KO (Fig. 3J). Most importantly and relevant to pro-inflammatory activation of macrophages, a total of 168 genes (70%) in Cluster-I are regulated by the Interferon-beta (IFN-β) pathway (Fig. 3I) and are therefore under the control of similar transcriptional regulation and factors. These results suggest that a reduction or complete deletion of Lamin-A/C significantly upregulates IFN-β related pro-inflammatory gene expression. Lamin-A/C reduction might drive the pro-inflammatory gene expression via IFN-β pathway, which we tested next.

### Augmentation of the IFN-β-STAT axis in LPS activated Lamin-A/C KO-BMDMs

To understand how *Lamin-A/C* reduction might regulate changes in pro-inflammatory gene expression, we subsequently focused on identifying associated transcription factors. Therefore, the promoters of 168 Lamin-A/C- and IFN-β-dependent genes were subject to a statistical overrepresentation analysis to detect enriched transcription factor binding sites. The analysis showed that the most overrepresented matrices corresponded to IRFs, p65, and STATs sites (Figs. 4A, S9A&B). This result gave a crucial hint that IRFs, NF-κB and STATs are potential transcription factors that might be influenced by *Lamin-A/C* deletion, which was probed in subsequent steps.

**Fig. 4:**
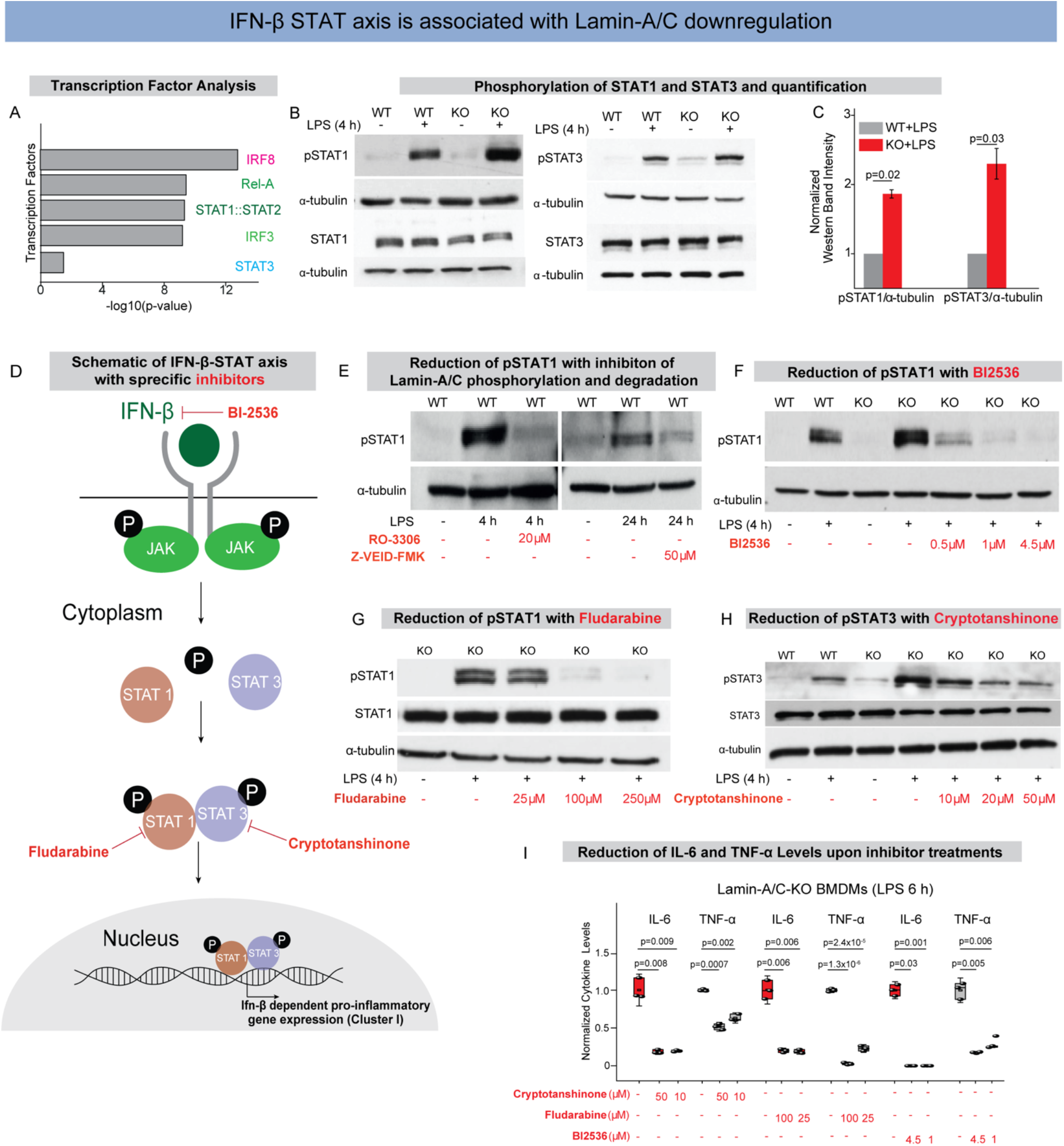
Degradation or knockout of Lamin-A/C augments the IFN-β-STAT axis during pro-inflammatory activation: **(A)** Bar graphs show transcription factors with corresponding -log10(p-value) found by statistical overrepresentation analysis to detect enriched transcription factor binding sites in the promoter of the 168 genes (LPS induced, and Lamin-A/C influenced Interferon-Beta dependent Genes). **(B)** Immunoblots show pSTAT1 (upper) and total-STAT1 (lower) **(Left)**, and pSTAT3 (upper), and total-STAT3 (lower) **(Right)** levels in wildtype (WT), WT+LPS, *Lamin-A/C* knockout (KO), and KO+LPS treated BMDMs. **(C)** Bar graphs show quantification of pSTAT1 and pSTAT3 normalized with α-tubulin. Calculated values were further normalized to the WT+LPS treated BMDMs condition over three independent experiments. Data are presented as Mean±S.E. **(D)** Scheme shows IFN-β-STAT axis and how the specific inhibitors used in this study inhibit phosphorylation of different STATs and thus inhibit pro-inflammatory gene expression. **(E)** Immunoblots show pSTAT1 levels in WT, WT+LPS treated BMDMs with RO-3306 and Z-VEID-FMK. **(F)** Immunoblot shows pSTAT1 levels in WT, WT+LPS, KO, KO+LPS and KO+LPS treated BMDMs with different concentrations of BI-2536. **(G)** Immunoblot shows pSTAT1 and total STAT1 levels in KO, KO+LPS and KO+LPS treated BMDMs with different concentrations of Fludarabine. **(H)** Immunoblot shows pSTAT3 and total STAT3 levels in WT, WT+LPS, KO, KO+LPS, and KO+LPS treated BMDMs with different concentrations of Cryptotanshinone. **(I)** Box plots show the lowering of secreted cytokine levels in LPS treated KO-BMDMs, treated with multiple concentration of Cryptotanshinone, Fludarabine, and BI-2536, as compared to only LPS treated KO-BMDMs. Levels were normalized to 6 h LPS treated BMDMs condition to find the fold change. In all the Immunoblot α-tubulin served as loading control. In all the box plots, boxes show 25th and 75th percentiles, the middle horizontal line shows the median, small open squares show the mean, and whiskers indicate S.D. *p*-values were obtained with the two-sided Student’s *t*-test. All experiments were independently repeated three or more times.

As a majority of the pro-inflammatory genes, whose upregulation correlates with Lamin-A/C reduction or loss, were predicted to be under the regulation of the STAT family of transcription factors (Figs. 4A, S9A&B), we first probed STAT1 and STAT3 (51). We could indeed confirm that the differential activity of STAT family of transcription factors underlies the observed upregulation of pro-inflammatory genes. As expected, levels of pSTAT1(Y701) and pSTAT3(Y705) are significantly increased in WT-BMDMs upon LPS treatment and that these levels were further augmented in LPS treated KO-BMDMs (Figs. 4B&C). Hence, STAT1/3 phosphorylation are modulated by Lamin-A/C reduction, but not the overall STAT1 or STAT3 levels (Fig. 4B). To probe whether Lamin-A/C reduction not only affects the phosphorylation status of STAT transcription factors, but also their recruitment to the promoter region of inflammatory genes, ChIP-qPCR experiments were conducted. Higher enrichment of pSTAT1 at the promoter regions of *IL-6* and *TNF-α* genes in LPS treated KO-BMDMs (Fig. S10A) confirmed higher transcriptional activity of pSTAT1 in LPS treated KO-BMDMs compared to WT-BMDMs, using IgG as control (Fig. S10B). To confirm that the STAT phosphorylation is indeed dependent on Lamin-A/C reduction, we show that inhibition of Lamin-A/C phosphorylation and degradation with the clinical pharmaceutical drugs RO-3306 and Z-VEID-FMK, respectively, could indeed reduce the LPS induced increase in STAT1&3 phosphorylation during macrophage inflammatory response (Fig. 4E). This gives first hints that the pro-inflammatory response of macrophages can indeed be reduced by targeting Lamin-A/C degradation.

To provide a potential mechanism of how the reduction of a nuclear envelope protein, i.e., Lamin-A/C might drive the phosphorylation of cytoplasmic proteins such as members of the STAT family, we checked the levels of Janus Kinase (JAK). Upon LPS activation, macrophages upregulate the expression levels of Type I interferons, such as interferon alpha and interferon beta (IFN-alpha and beta)(52)(Fig. S11A). Secreted IFNs bind their receptors and lead to the activation of signaling pathways, mainly Jak to activate STAT pathway (Figs. 4D&S11A). In response to IFNs binding to their receptors, Jaks get phosphorylated and in turn phosphorylate different STATs, mainly STAT1, 2 & 3, which then stimulate pro-inflammatory gene expression (52) (Figs. 4D&S11A). Thus, we checked whether reduction in Lamin-A/C levels influences expression levels of IFN-β or not. We confirmed a higher expression of *IFN-β* mRNA in LPS-treated KO BMDMs (Fig. S11B&C). At the same time, the expression of *IFN-β* mRNA depends on Lamin-A/C phosphorylation as WT BMDMs treated with LPS in the presence of RO-3306 showed significantly lower levels of *IFN-β* mRNA compared to only LPS treated BMDMs (Fig. S11D). This higher mRNA expression of *IFN-β* in KO BMDMs would lead to higher secretion of IFN-β and thus activation of IFN-β pathway. Indeed, we confirmed higher phosphorylation of Jak1 (Tyr 1034/1035) in LPS treated KO BMDMs as compared to LPS treated WT BMDMs (Fig. S12A). Altogether, this provides a plausible mechanism of how Lamin-A/C might regulate STAT activity via Janus Kinase.

To exclude that most amplified transcriptional responses in the KO-BMDMs does not reflect an increased hypersensitivity to LPS, but are primarily mediated by STAT1&3, we also checked the levels of other crucial transcription factor (STAT5, IRF3, and p65) activated during LPS treatment of macrophages, and found them to remained unchanged (Figs. S12B-D).

### Pharmaceutical inhibition of the IFN-β-STAT axis reverses the enhanced LPS response in Lamin-A/C KO-BMDMs

To test whether the enhanced LPS response in KO-BMDMs could be reversed by pharmaceutical inhibition of STAT pathway members, KO-BMDMs were treated with inhibitors to suppress the expression and activity of IFN-β and phosphorylation of STAT family of transcription factors. Indeed, different concentrations of BI2536, a specific drug to reduce IFN-β levels by inhibiting its transcription(53) and in clincal trials against various cancers(54, 55), are able to reduce the levels of pSTAT1 in LPS treated KO-BMDMs to similar levels as only LPS treated WT-BMDMs (Fig. 4F). BI-2536 inhibition also resulted in reduced mRNA levels of pro-inflammatory genes *IL-6* and *TNF-α* (Fig. S13), along with secreted IL-6 and TNF-α cytokines in LPS treated KO-BMDMs (Fig. 4I). To confirm this for STAT1, we used Fludarabine, a clinically approved specific inhibitor of pSTAT1(Y701)(56) used to address chronic lymphocytic leukemia (CLL)(57) and non-Hodgkins lymphoma (NHL)(58), which was able to counteract the hyper-upregulation of pSTAT1 levels in LPS treated KO-BMDMs (Fig. 4G). When specifically blocking STAT3 phosphorylation using the drug Cryptotanshinone(59), which is known for its anticancer activities(59), the normal activation levels of pSTAT3 in LPS treated KO-BMDMs were restored (Fig. 4H). Upon different drug treatments, levels of total STAT1 and STAT3 largely remained similar (Figs. 4G&H). Lastly, using these drugs in LPS treated KO-BMDMs, we could significantly reduce the levels of pro-inflammatory cytokines (IL-6 and TNF-α) in LPS treated KO-BMDMs (Fig. 4I). Altogether, these observations illustrate that indeed KO BMDMs have a hyper-active IFN-β pathway, and that IFN-β-STAT axis can be targeted by inhibiting Lamin-A/C phosphorylation and degradation.

### Inhibiting Lamin-A/C phosphorylation downregulates the inflammatory response of *E. coli* infected macrophages

Gram-negative bacteria, such as *E. coli*, activate macrophages due to the presence of LPS on their outer cell membrane. To validate that the mechanism described in this study is independent of whether macrophages get exposed to LPS in soluble form or exposed to gram-negative bacteria, we asked whether bacterial infection of macrophages also prompts the activation of inflammatory programs via the above described Lamin-A/C phosphorylation and degradation route. Therefore, BMDMs were exposed to heat-inactivated killed *E. coli* or active living *E. coli* and the expression status of *Lamin-A/C* was monitored 6 h post infection (Fig. 5A). Again, exposure to *E. coli* resulted in a significant reduction of *Lamin-A/C* mRNA levels within the first 6 h of exposure (Fig. 5B). This observation was accompanied by a significant Lamin-A/C degradation post 24 h heat-inactivated killed or active living *E. coli* infection (Fig. 5C). This confirms that LPS induced degradation of Lamin-A/C is inherent to macrophage activation by soluble LPS or cell wall bound LPS of gram-negative bacteria.

**Fig. 5:**
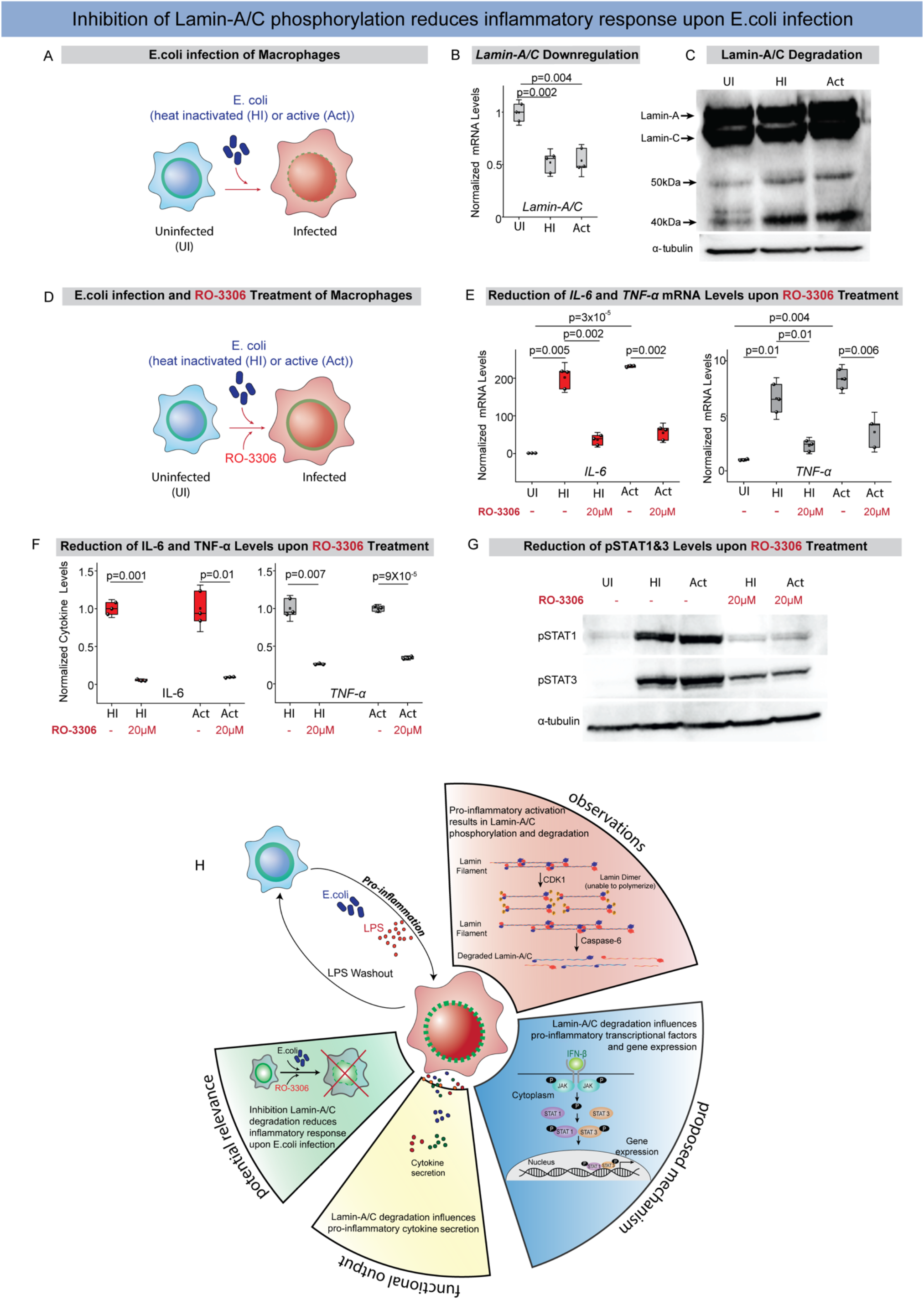
Inhibiting Lamin-A/C phosphorylation downregulates the inflammatory response of *E. coli* infected macrophages: **(A)** Scheme shows experimental setup-Wild Type BMDMs were infected with either heat inactivated (HI) or active (Act) *E. coli*. **(B)** Box plot shows mRNA levels of *Lamin-A/C* in uninfected (UI) or *E. coli* infected BMDMs as determined by qPCR. Levels were normalized to uninfected BMDMs condition to find the fold change. **(C)** Immunoblot shows Lamin-A/C degradation in *E. coli* infected BMDMs as compared to UI BMDMs. α-tubulin served as a loading control. **(D)** Scheme shows experimental setup-Wild Type BMDMs were infected with *E. coli* and treated with RO-3306. **(E)** Box plots show normalized mRNA levels of *IL-6* and *TNF-α* in untreated and E. coli infected BMDMs also treated with RO-3306. Values were further normalized to UI BMDMs condition to find the fold change. **(F)** Box plots show normalized secreted IL-6 and TNF-α cytokine levels in untreated and E. coli infected BMDMs also treated with RO-3306. Values were normalized to UI BMDMs condition to find the fold change. **(G)** Immunoblots show pSTAT1 and pSTAT3 in BMDMs infected with *E. coli* and treated with RO-3306 as compared to only *E. coli* infected BMDMs. α-tubulin served as loading control. **(H)** Schematic overview shows how pro-inflammatory activation by LPS or infection with *E. coli* of primary bone-marrow derived macrophages results in the phosphorylation and degradation of Lamin-A/C and its regulatory role in the pro-inflammatory gene expression and cytokine secretion via the IFN-β-STAT axis. In all the box plots, boxes show 25th and 75th percentiles, the middle horizontal line shows the median, small open squares show the mean, and whiskers indicate S.D. *p* values were obtained with the two-sided Student’s *t*-test. All experiments were independently repeated three or more times.

Towards therapeutic applications, we again asked whether the pro-inflammatory response of macrophages upon *E. coli* infection could also be scaled down by inhibiting Lamin-A/C phosphorylation with the pharmaceutical inhibitor RO-3306. Infecting BMDMs with heat-inactivated killed *E. coli*, or with active living *E. coli*, while inhibiting Lamin-A/C phosphorylation by RO-3306 pretreatment (Fig. 2H&5D), not only decreased the expression of pro-inflammatory genes (*IL-6*, and *TNF-*α) (Fig. 5E), but also the production and secretion of inflammatory cytokines (IL-6, and TNF-α) (Fig. 5F). We further also showed that *E. coli* induced phosphorylation of STAT1 and STAT3, as required for pro-inflammatory gene expression and cytokine synthesis, can be completely blocked by inhibiting Lamin-A/C phosphorylation and degradation (Fig. 5G). Taken together, we discovered here a new molecular route by which the cytokine production and secretion induced by gram-negative bacteria infections of macrophages might be reduced.

## Discussion and Outlook

As chronic inflammatory diseases are thought to be stimulated by the excessive secretion of pro-inflammatory cytokines and of other macrophage-derived metabolites, the discovery of additional potent regulators that control this excessive secretion is thus destined to initiate the research and development of novel therapeutic strategies and molecular targets. It had not been reported before that degradation of nuclear lamins are potential regulators of pro-inflammatory macrophage activation and of the subsequent secretion of inflammatory stimulants. Our data now suggest that Lamin-A/C phosphorylation and degradation is a prerequisite for the pro-inflammatory response of macrophages (Fig. 2). Inflammation could indeed be reduced by inhibiting Lamin-A/C degradation by either targeting CDK1 or Caspase-6 (Figs. 2&5), or by inhibiting the IFN-β-STAT axis, particularly pSTAT1 or pSTAT3 (Fig. 4). Several human diseases and pathological conditions, including muscular dystrophy and human inflammageing, have been shown to be associated with mutations or low-levels of Lamin-A/C along with higher levels of inflammatory metabolites circulating in the blood. Here, we discovered novel Lamin-A/C regulated events that modulate the pro-inflammatory response of macrophages, as discussed sequentially below (Fig. 5H).

First, we here discovered that macrophage activation and Lamin-A/C degradation are correlated (Fig. 1), and that Lamin-A/C phosphorylation by CDKs and its subsequent degradation by Caspase-6 controls pro-inflammatory gene expression and cytokine secretion (Fig. 2). Even though targeting CDKs has previously been shown to inhibit LPS induced activation of macrophages(60), the molecular events leading to inhibition were not known. Our pharmaceutical inhibition studies provide the regulatory mechanism showing that CDKs induced Lamin-A/C phosphorylation subsequently triggers Lamin-A/C protein degradation. Even though Lamin-A/C phosphorylation has recently been implicated in accelerated age associated pathologies like progeria(61),with potential connections to inflammatory gene expression, the actual role of Lamin-A/C has remained an unsolved mystery. Our data also shed light into another unsolved question, namely why Caspase-6 deficient mice show reduced TNF-α levels in plasma upon LPS-injection(62). This question was raised as Caspase-6 is classified as an effector or executioner caspase, based on its pro-apoptotic activity in cleaving other caspases and structural proteins like lamin(63–65). With our inhibitory experiments we now uncover the missing link suggesting that the degradation of Lamin-A/C by Caspase-6 controls pro-inflammatory gene expression and cytokine secretion.

Second, while the IFN-β-activated STAT family of transcription factors are known to drive the expression of pro-inflammatory genes, we discovered here that the Lamin-A/C phosphorylation and degradation upon pro-inflammatory activation of macrophages is necessary to augment the phosphorylation of STAT1 and STAT3, but not of STAT5 (Fig. 4). STAT1 and STAT3 are known to have functional and physical interactions and the expression of gene targets regulated by STAT1 and STAT3 is dependent on the relative proportions of STAT1 homodimers, STAT3 homodimers, and the STAT1/STAT3 heterodimers. Our observations do suggest higher activity of Janus-activated kinases, known kinases involved in STAT phosphorylation, yet whether Lamin-A/C degradation promotes their heterodimer formation remains unknown. Thus, how mechanistically Lamin-A/C, a nuclear protein, achieves the regulation of a kinase that acts at the cell membrane levels still needs to be further uncovered. Only recently, it has been reported that Emerin, another nuclear envelope protein bound to Lamin-A/C, binds STAT3 and by this has an impact on STAT3 retention to the nuclear membrane(66). Since Lamin-A/C loss or degradation affects Emerin localization(50), we hypothesized that Lamin-A/C might influence STAT3 activity via this route, however further studies will need to be conducted to understand how other nuclear envelope proteins might regulate the JAK-STAT axis during the pro-inflammatory response. With the identified key players here, we hope that our findings stimulate many further explorations of the underpinning mechanisms and therapeutic innovations.

Finally, data obtained from our study advocate the efficacy of targeting nuclear lamina proteins and associated molecular mechanism to protect macrophages from virulent *E. coli* infection-induced inflammation. Moreover, the ability of these targets to tame the pro-inflammatory macrophage response could be attributed to their ability to downregulate the TLR4 and STAT pathways (Fig. 5). Therefore, targeting nuclear lamina could be an ideal candidate immunomodulator and anti-inflammatory agent against diseases caused by *E. coli* infection. However, further studies using animal infection models will be needed to warrant the *ex vivo* and *in vivo* efficacy of these targets. Especially, the development of animal infection models using other pathogens than *E. coli* would further broaden our discovery, as already in tuberculosis-infected mice *Lamin-A/C* mRNA levels are significantly downregulated in alveolar macrophages(67).

Towards the clinical significance of our findings, we confirmed that several different tissue resident macrophages, and not only BMDMs, exhibit reduced *Lamin-A/C* mRNA levels in inflamed tissues (Fig. 1), suggesting a major yet generalized role of the nuclear lamina during macrophage activation. These findings indicate that the nuclear lamina and associated proteins might be able to serve as novel therapeutic targets to reduce inflammation in multiple tissues across several chronic inflammatory diseases. The generalization of these findings, showing that inflammation finally results in a reduced *Lamin-A/C* expression also in other primary macrophages of rodents, humans, and other species (Fig. 1) is essential for the exploitation of this knowledge for clinical applications. We also demonstrate that the observed dependencies can be observed not only upon LPS stimulation, but that the activation of macrophages as induced by virulent gram-negative bacteria, here upon *E. coli* infection, also leads to a degradation of Lamin-A/C (Fig. 5), and importantly that the pro-inflammatory response of macrophages can be reduced by targeting the enzymes involved (Fig. 5). Exploiting the ability of these targets to downregulate the STAT pathways (Fig. 4) provides new opportunities to develop immunomodulator and anti-inflammatory agents to cope with acute bacterial infections, or chronic inflammation. This study provides mechanistic insights into how macrophage inflammatory response during several pathological conditions could potentially be controlled by therapeutically targeting molecular intermediates involved in Lamin-A/C phosphorylation (CDKs), proteases necessary for Lamin-A/C degradation (Caspase-6) and in the IFN-β-STAT expression axis. Using different pharmacological inhibitors to inhibit Lamin-A/C phosphorylation, degradation, and phosphorylation of STAT1/3 within the IFN-β-STAT-axis, we have identified possible molecular targets in proof-of-concept experiments. However, further studies using animal infection models will be needed to warrant the ex vivo efficacy of these targets. In support of the clinical significance, it is highly relevant that tissue resident macrophages in a variety of inflamed human tissues show a *Lamin-A/C* mRNA downregulation, concomitant with an upregulation of pro-inflammatory genes (Fig. 1). Even though these targets do influence the cell-cycle, a major consortium has recently started a phase II trial to target CDKs as an adjunctive therapy in rheumatoid arthritis, suggesting the possibility of targeting CDKs and Caspases as a potential therapeutic against inflammation. Lastly, our work also opens novel avenues to explore the potential role of other nuclear lamina related proteins in taming pro-inflammatory reactions and the far less well-understood processes of resolving inflammatory episodes.

## Acknowledgements

The authors thank the Mice Facility at ETH Zurich and Prof. Annette Oxenius (ETH Zurich) for donating post-mortem animals for bone marrow isolation and Chantel Spencer for technical support in experiments like ELISA. The authors thank Prof. Manfred Kopf (ETH Zurich) for providing alveolar macrophages. We also want to thank Cornell University Institutional Animal Care and Use Committee and Hannah Fong for isolating and sending bone-marrow from KO mice. We further thank the ScopeM facility at ETH Zurich for access to confocal microscopy and the Genetic Diversity Center at ETH. This work was supported by ETH Zurich, SNF NCCR ‘Molecular Systems Engineering, SNF CR32I3_156931, an ETH Research Grant ETH-24 18-1 grant to V.V., the National Institutes of Health (NIH) awards R01HL082792, R01GM137605, and U54CA210184 to J.L., University of Birmingham (UoB), and SNF SPARK award CRSK-3_195952 to N.J.

## Author contributions

NJ conceived of the project. JM and NJ designed and performed the experiments. JM, and NJ analyzed the data. AE isolated and provided the bone-marrow isolated from Lamin-A/C-KO mice and JL, and MM accompanied the work through discussions. JM, JL, VV, and NJ wrote the manuscript. All authors edited and approved the manuscript.

## Materials and Methods

Animal: Lamin-A KO (Lamin-A-/-) have been described previously(50). Lamin-A mutant mice were provided with gel diet supplement (Nutri-Gel Diet, BioServe) to improve hydration and metabolism following the onset of phenotypic decline. Maintenance and euthanasia of animals were performed in accordance with relevant guidelines and ethical regulations approved by the Cornell University Institutional Animal Care and Use Committee, protocol nos. 2011-0099 and 2012-0115.

### Bone marrow isolation, cell culture and primary macrophage differentiation

Femurs were isolated from post-mortem healthy mice (5–7 weeks old C57BL/6) and bone marrow was flushed with PBS. Bone marrow was further passed through a 7 μm cell strainer to obtain a single-cell suspension. Bone marrow was centrifuged and suspended in BMDM medium. Four million cells were seeded in 60 mm nontreated plastic dishes for 7 days in 5% CO_2_ at 37 °C. Equal volumes of medium were again added on day 4. BMDMs were used on day 7 for further experiments. BMDM culture medium contained Dulbecco’s modified Eagle’s medium (DMEM) (Cat No. 31966-021, Gibco), 10% (v/v) fetal bovine serum (FBS), glutamine (Cat No. 35050-038, Gibco), non-essential amino acids (Cat No. 11140-035, Gibco), sodium pyruvate (Cat No. 11360-039, Gibco), β-mercaptoethanol (Cat No. 31350-010, Gibco), penicillin–streptomycin (Cat No. 15140-122, Gibco) and L929 medium. L929 medium was prepared by culturing 2 million L929 cells in 200 ml medium in a 300 cm^2^ flask for 8 days without changing the medium. The medium used for L929 cell culture contained RPMI, 10% FBS, glutamine, nonessential amino acids, sodium pyruvate, HEPES, and β-mercaptoethanol. L929 fibroblast cells were cultured and maintained in 5% CO_2_ at 37 °C.

### Alveolar macrophage isolation

Alveolar macrophage isolation: Mice were euthanized by overdose (400 mg/kg body weight) of sodium pentobarbital by i.p. injection. Following anesthesia, a tracheostomy tube was placed, and mouse lungs were washed four times with 10 ml of sterile cold PBS (pH 7.4) containing 2 mM EDTA. The recovered Bronchoalveolar lavage was centrifuged at 1500 rpm for 7 min and the cells were harvested. The cell pellet was then resuspended in sterile medium and cultured for 2 hours before activating with LPS.

### RAW264.7 macrophage cell culture

RAW264.7 macrophages were cultured in DMEM (Cat No. 31966-021, Gibco), containing 10% (v/v) FBS and 1% penicillin–streptomycin (Cat No. 15140-122, Gibco) at 37 °C, 5% CO_2_.

### Macrophage activation

All studies were performed with primary BMDMs unless stated differently. Unlike the frequently used immortalized cell lines, BMDMs more closely mimic macrophage behaviour in vivo. Pro-inflammatory activation was achieved by stimulating macrophages with LPS (50 ng/ml) (L3129 Sigma). TNF-α treatment was done by treating BMDMs with 100ng/ml TNF-α (410-MT-010, R&D Systems).

### Drug treatments

Phosphorylation of STAT1 was inhibited by treating BMDMs with BI-2536 (1129 Axon Medchem). BMDMs were treated with different concentration of BI-2536 for 1 h before activating with LPS. BI-2536 was kept in the culture-media during LPS treatment. Phosphorylation of STAT3 was inhibited by treating BMDMs with Cryptotanshinone. BMDMs were treated with different concentration of Cryptotanshinone for 2 h before activating with LPS. Cryptotanshinone (16987 Cayman Chemical) was kept in the culture-media during LPS treatment. For Fludarabine (3495 R&D Systems) treatment, BMDMs were treated with different concentration of drug for 24 h before activating with LPS and Fludarabine was kept in the culture-media during LPS treatment. Phosphorylation of Lamin-A/C was inhibited with RO-3306 (AG-CR1-3515-M005 AdipoGen). BMDMs were treated with different concentration of RO-3306 for 2 h before activating with LPS. RO-3306 was kept in the culture-media during LPS treatment (50 ng/ml) (L3129 Sigma). Degradation of Lamin-A/C was inhibited by Caspase-6 Inhibitor Z-VEID-FMK (FMK006 R&D Systems). BMDMs were treated with different concentration of Caspase-6 Inhibitor Z-VEID-FMK for 2 h before activating with LPS. Caspase-6 Inhibitor Z-VEID-FMK was kept in the culture-media during LPS treatment. LB-100 (S7537 Lubio Science) was used to inhibit PP2A activity. BMDMs were treated with 2.5μM LB-100 for 2h before activating with LPS. LB-100 was kept in the culture-media during LPS-treatment (50 ng/ml) (L3129 Sigma).

### Immunostaining

BMDMs were rinsed twice with 1X PBS, followed by fixation using 4% paraformaldehyde (18814-20, Polysciences Inc.) in 1X PBS for 20 min. BMDMs were washed and permeabilized with 0.3% Triton X-100 (Sigma) in 1X PBS for 10 min. After washing twice, BMDMs were treated with 2% BSA (blocking solution) for 60 min before incubating with primary antibody (diluted in 2% BSA) at 4 °C overnight. Primary antibodies against Lamin-A/C (1:100, ab8980 Abcam; 1:100, ab8984 Abcam; 1:100 4777S Cell Signalling), phospho-Ser22-Lamin-A/C (pSer22) (1:100, 2026 Cell Signalling) and p65 (NFκB p65 1:300, sc-8008 Santa Cruz) were used for staining. BMDMs were then washed with blocking solution and incubated with the corresponding secondary antibodies, diluted in blocking solution, along with the nuclear stain Hoechst-33342 (1:1000, 62249 Life Technologies) for 45 min. Filamentous actin (F-actin) was labelled using phalloidin Alexa-Fluor 568 (1:200, Life Technologies). All antibodies used in this study have previously been extensively used and validated for immunofluorescence studies.

### Confocal imaging and analysis

Images of single BMDM were taken using a Leica SP8 confocal microscope. To estimate spreading areas, 2D images were taken using a 40X (0.7 NA) air objective. To estimate protein expression levels, BMDMs were stained with the desired antibodies and 3D confocal images were captured rather than projected images. Fluorescence images were captured using a 63X (1.25 NA) oil objective with a z-step of 0.5 μm and a confocal pinhole of 1 A.U. Imaging conditions were kept the same during all experiments. All the quantifications per cell were carried out using MATLAB2019b (MathWorks) or ImageJ (NIH). Imaging conditions were kept the same during all experiments.

### Cytokine secretion

Cell-culture medium was collected post drug treatment or LPS stimulation for desired time (mentioned in the respective figures). To remove cell debris, medium was centrifuged, and the supernatant was stored at -80 °C till further processing. ELISA kits were purchased from Peprotech TNF-α (900-TM54), and IL-6 (900-TM50) and experiment was performed according to the manufacturer’s protocol. All data fitting and quantifications were done using OriginPro8.1 and MATLAB2019b (MathWorks), respectively.

### Immunoblotting

Briefly, BMDMs were washed once with ice-cold PBS and lysed for 20 min with lysis buffer (20mM TRIS,150mM NaCl, 1mM EGTA, 1mM EDTA, 1% Triton-X-100) supplemented with phosphatase inhibitor (1:100 P5726, Sigma Aldrich) and protease inhibitor (1:10 11836170001, Sigma-Aldrich) on ice. BMDMs were scratched and lysate was collected, homogenized and centrifuged at 4°C. Protein concentrations were determined with a BCA protein assay kit (23225, Thermo Fischer Scientific) and cell lysates were incubated with 5X Reducing Sample Buffer (0.3M Tris-HCL pH6.8, 100mM DTT, 5% SDS, 50% Glycerol, Pyronin G) for 5 minutes at 95°C. Protein lysate was separated using 10% Bis-Tris polyacrylamide gel. Proteins were transferred by wet tank transfer (110V 1hour) on to a nitrocellulose membrane. Membranes were blocked for 1 h using 5% BSA in TBS-T (0.01% Triton-X) and incubated overnight with primary antibodies (1:500 pSTAT1, 7649 Cell Signaling; 1:500 pSTAT3, 9145 Cell Signaling; 1:500 pSTAT5, 9359S Cell Signaling; 1:500 STAT1, 9172 Cell Signaling; 1:500 STAT3, 4904 Cell Signaling; 1:500 STAT5, 94205 Cell Signaling; 1:100 pJAK1, 3331S Cell Signaling; 1:250 pIRF3, 4947 Cell Signaling; 1:100 Lamin-A/C, 4777S Cell Signalling; 1:100 p-Lamin-A/C (Ser22), 2026 Cell Signalling; 1:1000 alpha-tubulin, 15246 Abcam; 1:100 pCdk1, 194874 Abcam) in 5% BSA TBST (0.01% Triton-X). Membranes were washed, incubated for 1 h with 1:10000 (donkey anti-rabbit secondary HRP Jackson or respectively donkey anti-mouse secondary HRP Jackson) in 5% BSA TBS-T (0.01% Triton-X) and washed again before exposing to Immunoblotting Substrate (Pierce ECL Immunoblotting Substrate 32209 Thermo Fisher Scientific).

### Quantification of immunoblots

The films were scanned and converted into 8-bit (grayscale) images. For each protein of interest single regions of interest were defined and the mean gray value was measured for the protein and for the background. To get relative amounts as a ratio of each protein band relative to the lane’s loading control, single regions were as well measured for the single bands of the loading control and the background of the loading control and protein of interest. The mean gray values were inverted (255-mean gray value) and the inverted background was subtracted from the inverted protein value to get the net protein value. The net protein value was divided by the net loading control value to get relative amounts of the protein of interest.

### RNA isolation and Real-Time qPCR analysis

RNA was isolated with a Qiagen RNA Isolation Kit according to the manufacturer’s protocol. cDNA was prepared using iScript Advanced cDNA Synthesis Kit (172-5038 BioRad). Real-time qPCR experiments were performed using SsoAdvanced™ Universal SYBR® Green Supermix (1725271 BioRad) in a CFX96 Real-Time PCR Detection System (Bio-Rad). Primers are listed in the following:

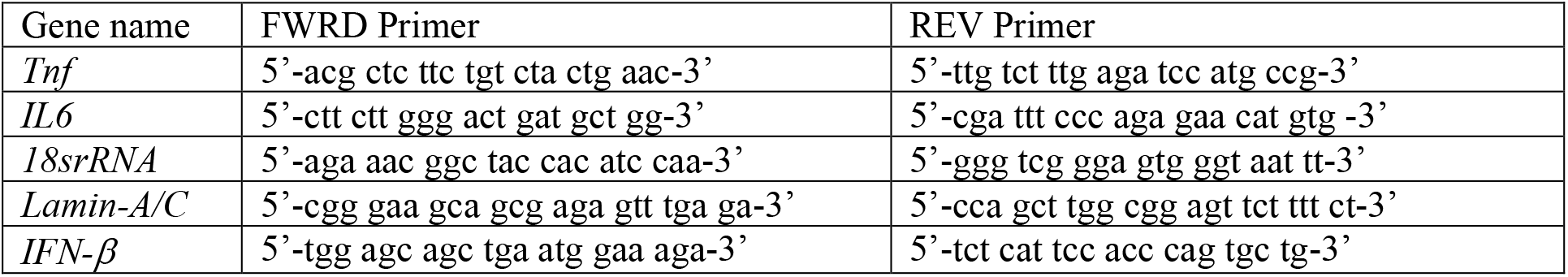

The qPCR analysis is explained here with exemplary (triplicate average) Ct values produced during real-time q-PCR analysis:

|  | Raw Ct values 18s rRNA | Raw Ct values gene of interest |
| --- | --- | --- |
| WT | 6.59 | 26.42 |
| WT+LPS | 7.22 | 17.84 |

<u>Step 1: Normalization of Ct-values</u>

Raw Ct values (18s rRNA) were subtracted of the Raw Ct values (gene of interest) of the sample.

Step 1: Normalization of Ct-values Raw Ct values (18s rRNA) were subtracted of the Raw Ct values (gene of interest) of the sample.
|  | Normalized Raw Ct values |
| --- | --- |
| WT | 26.42-6.59=19.83 |
| WT+LPS | 17.84-7.22=10.62 |

<u>Step 2: Normalization to WT sample</u>

|  | Normalized Ct values |
| --- | --- |
| WT | 19.83-19.83=0 |
| WT+LPS | 10.62-19.83=-9.21 |

<u>Step 3: Calculation of Fold-Change</u>

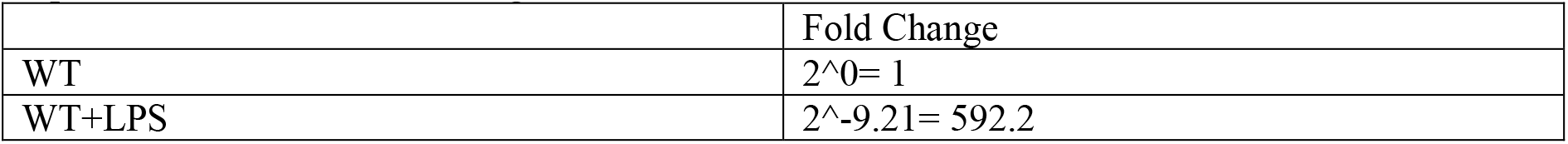

|  | Fold Change |
| --- | --- |
| WT | 2<sup>0</sup> = 1 |
| WT+LPS | 2<sup>-9.21</sup> = 592.2 |

In this case the calculated fold change would be 592.2 when comparing the gene expression between the WT and WT+LPS treated BMDMs samples. This calculated fold change was then plotted in Origin Pro.

### RNA Sequencing

RNA was isolated with the Qiagen RNA Isolation Kit according to the manufacturer’s protocol. For library preparation, the quantity and quality of the isolated RNA were determined with a Qubit (1.0) Fluorometer (Life Technologies) and a Tapestation 4200 (Agilent). The TruSeq Stranded HT mRNA Sample Prep Kit (Illumina) was used in subsequent steps following the manufacturer’s protocol. Briefly, total RNA samples (1 μg) were ribosome-depleted and then reverse-transcribed into double-stranded cDNA with actinomycin added during the first-strand synthesis. The cDNA samples were fragmented, end-repaired and polyadenylated before ligation of TruSeq adapters. The adapters contain the index for dual multiplexing. Fragments containing TruSeq adapters on both ends were selectively enriched with PCR. The quality and quantity of the enriched libraries were validated using a Qubit (1.0) fluorometer and the Tapestation 4200 (Agilent). The product is a smear with an average fragment size of ∼360 bp. The libraries were normalized to 10 nM in Tris-Cl 10 mM, pH 8.5 with 0.1% Tween 20. For cluster generation and sequencing, a Hiseq 4000 SR Cluster Kit (Illumina) was used using 8 pM of pooled normalized libraries on the cBOT V2. Sequencing was performed on the Illumina HiSeq with single-end 125 bp using the HiSeq 3000/4000 SBS Kit (Illumina). Reads quality was checked with FastQC (https://www.bioinformatics. babraham.ac.uk/projects/fastqc/) and sequencing adapters were trimmed using Trimmomatic63. Reads at least 20 bases long, and an overall average Phred quality score greater than 10 were aligned to the reference genome and transcriptome (FASTA and GTF files, respectively, downloaded from Ensemble, genome build GRCm38) with STAR v2.5.164, with default settings for single end reads. The distribution of reads across genomic features was quantified using the R package Genomic Ranges from Bioconductor Version 3.065. Differentially expressed genes were identified using the R package edgeR from Bioconductor Version 3.066.

### Functional analysis of genes

Gene Ontology and KEGG pathway analysis of differentially regulated genes was performed using DAVID software. Prediction of Transcription factors and their binding analysis was performed using PSCAN and Interferome.

### Reanalysis of previously published RNA-Sequencing data

The data downloaded from the public repositories was formatted in different manners. To give an insight of our analysis we will explain how our analysis was done based on one example. In the following, we will explain how we extracted the data for murine microglia treated with LPS for 6 hours (GSE90046).

#### Step 1 Downloading and Filtering the data

We downloaded the respective data from the GEO repository (GSE90046) and subsequently filtered the data for the genes of interest (*Lmna* and pro-inflammatory genes). The Sequencing data was given, in this case, in counts per gene (in triplicates).

#### Step 2 Normalizing data to total counts

To normalize with the total counts, we calculated to number of total counts of the RNA-Sequencing per sample (by adding up all counts across the sample) and the divided the respective counts of our gene of interest by the total number of counts.

#### Step 3 Calculating the log2fold-change

Subsequently we took the mean of the triplicates (of the gene of interest) and calculated the log2fold change comparing untreated and 6 h LPS treated murine microglia. This was done by making the ratio of the means (Mean fragments of Control samples/ Mean fragments of Control 6 h LPS samples) and then calculating the log2 of the resulting ratio.

#### Step 3 Plotting the data

To plot the log2fold changes in one big panel as done in Fig. 1A, we gathered all the log2fold-changes of the different species and cell types and plotted these in a heatmap by using Matlab2020.

### ChIP-Qpcr

10 million BMDMs per condition were cultured and treated accordingly. After 6 h of LPS treatment 16% Formaldehyde (18814-20, Polysciences Inc.) was added to the medium in the dish to a concentration of 1% and incubated (gently mixing) for 10 min. 1 ml of ice-cold 1.25 M glycine was added and incubated (gently mixing) for 5 min. The medium was discarded, and the dishes transferred onto ice. BMDMs on the dishes were washed with 5 ml of ice-cold PBS supplemented with protease inhibitor and phosphatase inhibitor (1:10 11836170001, Sigma-Aldrich, 1:100 P5726, Sigma Aldrich – PBS/c) and scratched in 2 ml PBS/c, which was collected. The dishes were washed with 1ml PBS/c and pooled with the 2 ml. Scratched BMDMs were centrifuged for 8 min at 300 g and 4 °C. The supernatant was aspirated, and the dry pellet snap frozen on dry ice. The pellet was thawed on ice and resuspended in 50 μl SLB/c buffer (10 mM Tris-HCL pH 8, 10 mM EDTA, 1% SDS supplemented with protease inhibitor (11836170001, Sigma-Aldrich)). Chromatin was sheared with a Covaris S220 AFA System (Covaris) with the following settings (Peak Power 140, Duty Factor 10.0, Cycles Burst 2000, Repetitions 15, 30 s ON/ 30 s OFF) at 4°C. Samples were centrifuged for 5 min at 20000 rpm and 4 °C and the supernatant transferred into low-binding tubes. 9 volumes (450 μl) of CDB/c (16.7 mM Tris-HCl pH8, 167 mM NaCl, 1.2 mM EDTA, 1.1% Triton X-100, 0.01% SDS supplemented with protease inhibitor and phosphatase inhibitor (1:10 11836170001, Sigma-Aldrich, 1:100 P5726, Sigma Aldrich) was added to the sample and incubated for at least 30 min with 25 μl Dynabeads A (10001D, Thermo Fischer Scientific) resuspended in CDB/c. After incubation the supernatant was transferred and 1% of the volume was stored separately at -20 °C (input). The remaining sample was incubated o/n on a rotary shaker at 4 °C with 2 μg of the appropriate antibody (IgG, 12-370 Merck; pSTAT1, 7649 Cell Signaling). The next day 25 μl Dynabeads resuspended in CDB/c were added and incubated for at least 1 h at 4 °C on a rotary shaker. Beads were washed sequentially with low-salt wash buffer (20 mM Tris-Hcl pH 8, 150 mM NaCl, 2 mM EDTA, 1% Triton X-100, 0.1% SDS), high-salt wash buffer (20 mM Tris-HCl pH 8, 500 mM NaCl, 2 mM EDTA, 1% Triton-X-100, 0.1% SDS), LiCl wash buffer (10 mM Tris-HCl pH 8, 250 mM LiCl, 1 mM EDTA, 1% sodium deoxycholate, 1% Igepal-CA630) and TE (10 mM Tris-HCl pH8, 1 mM EDTA). Immuno-precipitants were eluted from beads with Elution Buffer (100 mM NaHCO_3_, 1% SDS) incubating 15 minutes on a rotary shaker. 20 μl of 5 M NaCl was added to the elutions and to the with EB diluted 1% input sample and incubated at 65 °C and 95 rpm for 2 h. 10 μl o f 0.5 M EDTA, 20 μl 1 M Tris-HCl pH 6.5 and 1 μl of 20 mg/ml proteinase K was added to each sample and incubated for 1 h at 45 °C. DNA was extracted with the phenol chloroform extraction protocol. After DNA extraction the DNA-concentrations of the samples were measured. DNA samples were diluted, and the immunoprecipitation reaction assessed by qPCR. Primers are listed in the following:

| Gene name | FWRD Primer | REV Primer |
| --- | --- | --- |
| *Tnf-α* | 5'- tcc tga ttg gcc cca gat tg -3' | 5'- tag tgg ccc tac acc tct gt-3' |
| *IL6* | 5'- ccc acc ctc caa caa aga tt -3' | 5'- gct cca gag cag aat gag cta -3' |

The ChIP fold enrichments were calculated with the % Input–Method. This method is described in the following:

The signals obtained from ChIP are divided by signals obtained from an Input sample. This Input sample represents the amount of chromatin used in the ChIP. In these experiments a 1% of starting chromatin was used as Input. The calculation was done as following, example is illustrated below:

<u>Step 1: Adjusting the Input</u>

|  |  | Adjusted input to 100% |
| --- | --- | --- |
|  | Raw Ct | Ct Input – 6.644 |
| Input 1% (WT+LPS sample) | 32.7 | 26.1 |

<u>Step 2: Percent Input calculation</u>

<u>With the Adjusted Input (26.1) calculated the Percent Input of the sample was calculated.</u>

Step 2: Percent Input calculation With the Adjusted Input (26.1) calculated the Percent Input of the sample was calculated.
|  |  | Percent Input |
| --- | --- | --- |
| | Triplicate average Ct | $100 \cdot 2^{(26.1 - \text{Ct(IP)})}$ |
| pSTAT1 (WT+LPS sample) | 31.3 | 2.7 |

<u>These calculations were done for the different samples with their respective Input samples.</u>

<u>Step 3: Fold Change calculation</u>

To calculate the fold change enrichment the single samples were divided by the calculated Percent Input of the WT+LPS sample (2.7).

|  | Percent Input | Fold Change enrichment (Percent Input/2.7) |
| --- | --- | --- |
| pSTAT1 (WT+LPS sample) | 2.7 | 1 |
| pSTAT1 (KO+LPS sample) | 7.2 | 2.6 |

This calculated fold change enrichment was subsequently plotted in OriginPro.

### Bacterial infection

For active *E. coli* infection, bacterial strains MG1655 was used(68). For heat inactivated E. coli infection, bacterial strain ATCC35218 was used(69). *E. coli* strains were grown in LB medium and diluted in antibiotic-free BMDM culture medium before experiments after overnight growth to an optical density of 0.02. For heat inactivated *E. coli* infection, heat inactivated *E. coli* were added to the BMDMs in a concentration of 2.5×10^6^ bacteria/ml.

### Statistical analysis

The cell number assayed for quantification is given in individual figures. Comparisons were performed by means of a two-tailed Student t-test. All the statistic and data plotting were done in OriginPro software.

### Data and materials availability

All data, code, and materials used in the analysis will be made available for purposes of reproducing or extending the analysis.

