## Supplementary Data File for "Blockage of Lamin-A/C loss diminishes the pro-inflammatory macrophage response"

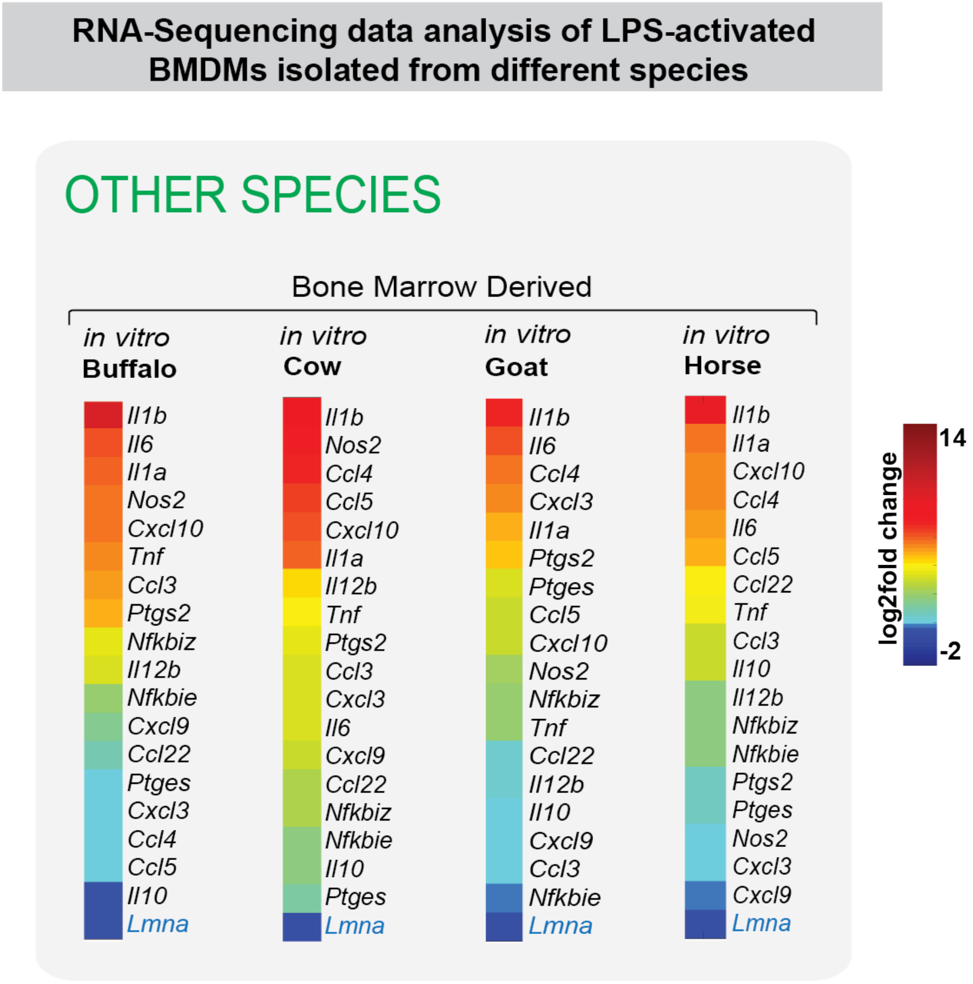


**Fig. S1: RNA-Sequencing data analysis of different species shows downregulation of *Lamin-A/C* mRNA expression levels in LPS-activated macrophages:** Color coded map shows the mRNA expression levels of various pro-inflammatory genes and *Lamin-A/C* in bone marrow derived macrophages isolated from other species (buffalo, cow, goat and horse) and treated with LPS. Expression data were obtained from public repositories(29).

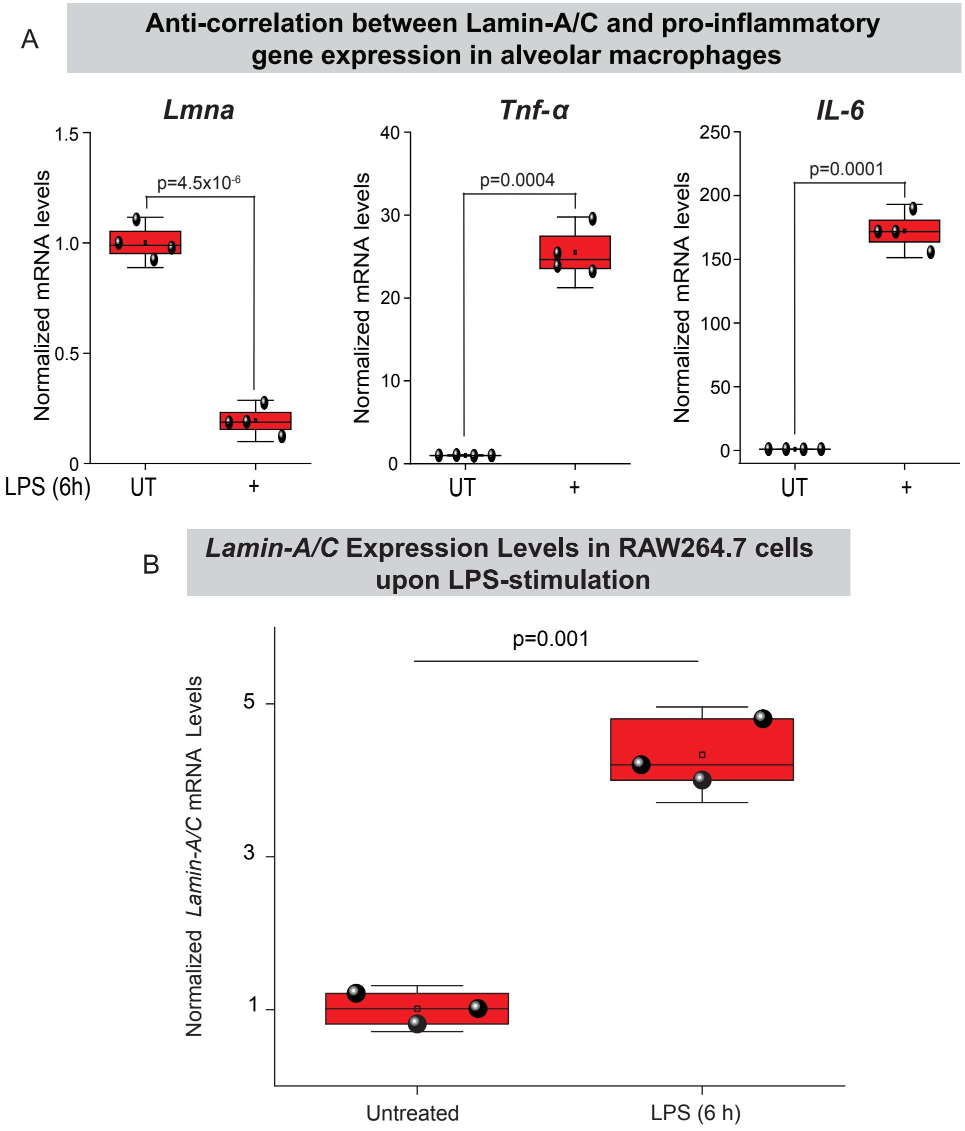


**Fig. S2: *Lamin-A/C* mRNA expression levels are downregulated in alveolar macrophages, while they are upregulated in RAW264.7 macrophages: (A)** Box plots show levels of *Lamin-A/C*, *Tnf-α* and *IL-6* mRNA in Untreated (UT) and 6 h LPS treated alveolar macrophages. Levels were normalized to UT alveolar macrophages condition to find the fold change. **(B)** Box plots show levels of *Lamin-A/C* mRNA in untreated and 6 h LPS treated RAW264.7 macrophages. Levels were normalized to UT RAW264.7 macrophages condition to find the fold change. In all the plots, the boxes show 25th and 75th percentiles, the middle horizontal line shows the median, small open squares show the mean, and whiskers indicate S.D. *p* values were obtained with the two-sided Student´s *t*-test. All the experiments were independently repeated three or more times.

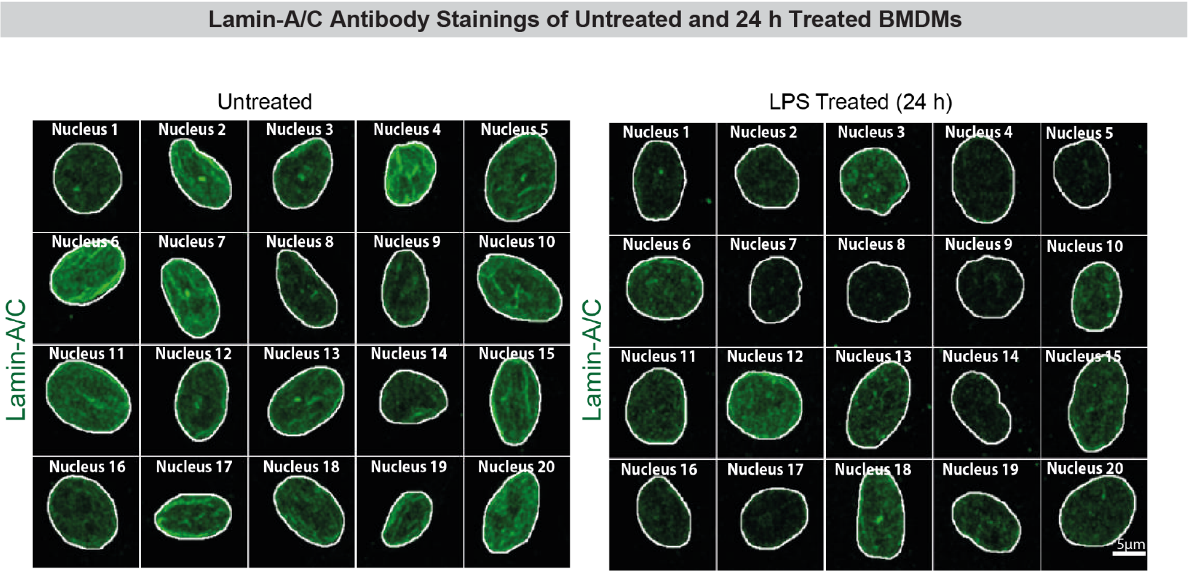


**Fig. S3: Collage of 20 macrophage nuclei stained for Lamin-A/C in Untreated and 24 hours LPS treated BMDMs. Scale bar = 5 μm.**

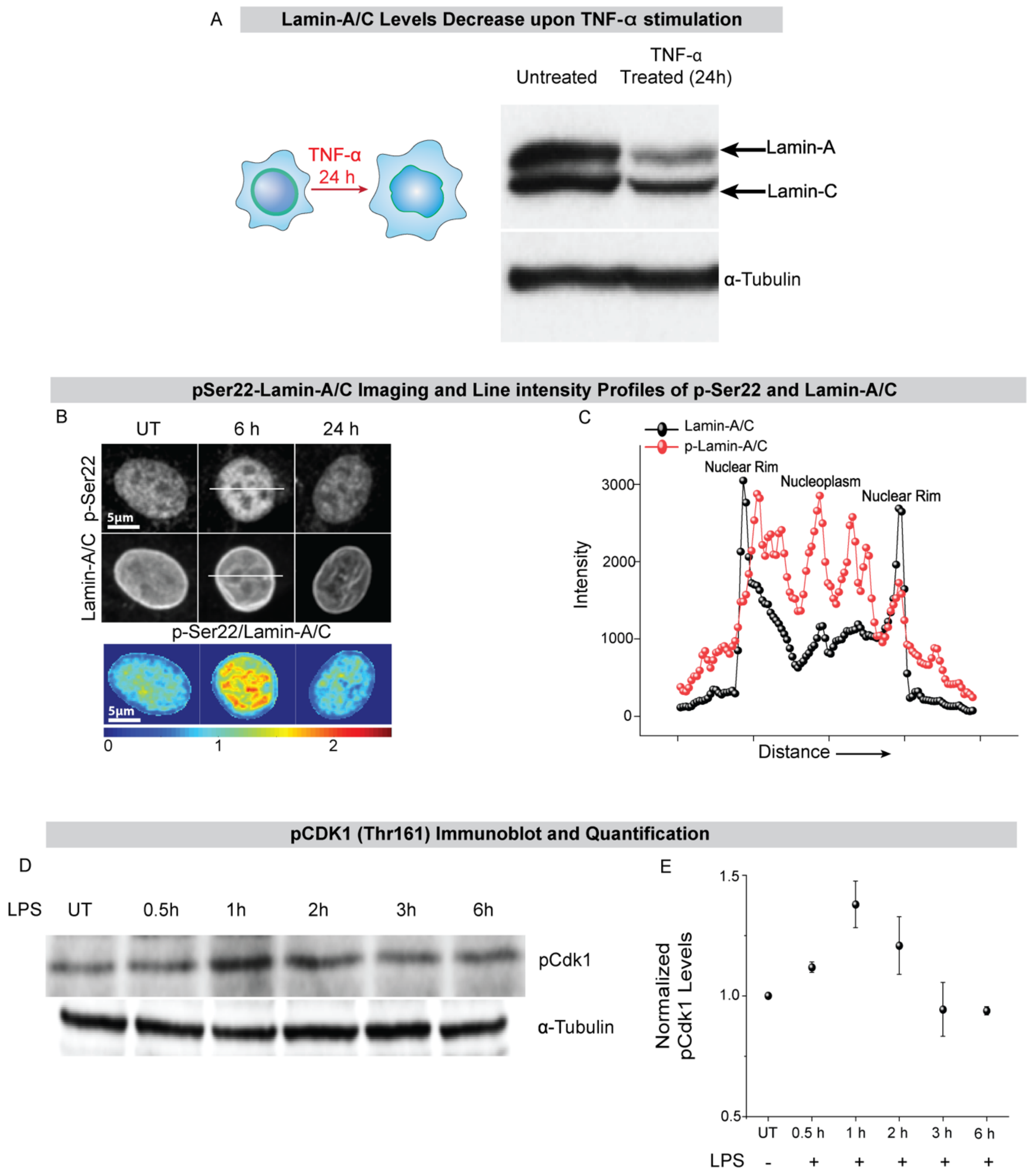


**Fig. S4: Quantification of Lamin-A/C protein levels upon TNF-α stimulation, and pSer22-Lamin-A/C and CDK1 (Thr161) levels upon LPS treatment: (A)** Immunoblot shows total levels of Lamin-A/C in Untreated (UT), and 24 h TNF-α treated BMDMs. α-tubulin served as a loading control. **(B)** Representative images of UT, 6 h and 24 h LPS treated BMDM stained with p-Ser22-Lamin-A/C and total Lamin-A/C antibodies. Scale bar = 5 µm. Also shown are the color-coded images showing the ratio of levels of p-Ser22-Lamin-A/C to total Lamin-A/C levels in the same nucleus. Scale bar = 5 µm. **(C)** Line intensity profiles (corresponding to white lines in B) of Lamin-A/C (in black) and p-Ser-22 Lamin-A/C (in red) in 6 h LPS BMDMs. **(D)** Immunoblots show levels of p-(Thr161)-CDK1 in BMDMs treated with LPS for different periods of time. α-tubulin served as a loading control **(E)** Time course quantification of normalized p-CDK1 protein levels. p-CDK1 levels were normalized to α-tubulin. Calculated values were normalized to the UT BMDMs condition over three independent immunoblotting experiments. Data are presented as Mean±S.E. All the other experiments were independently repeated three or more times

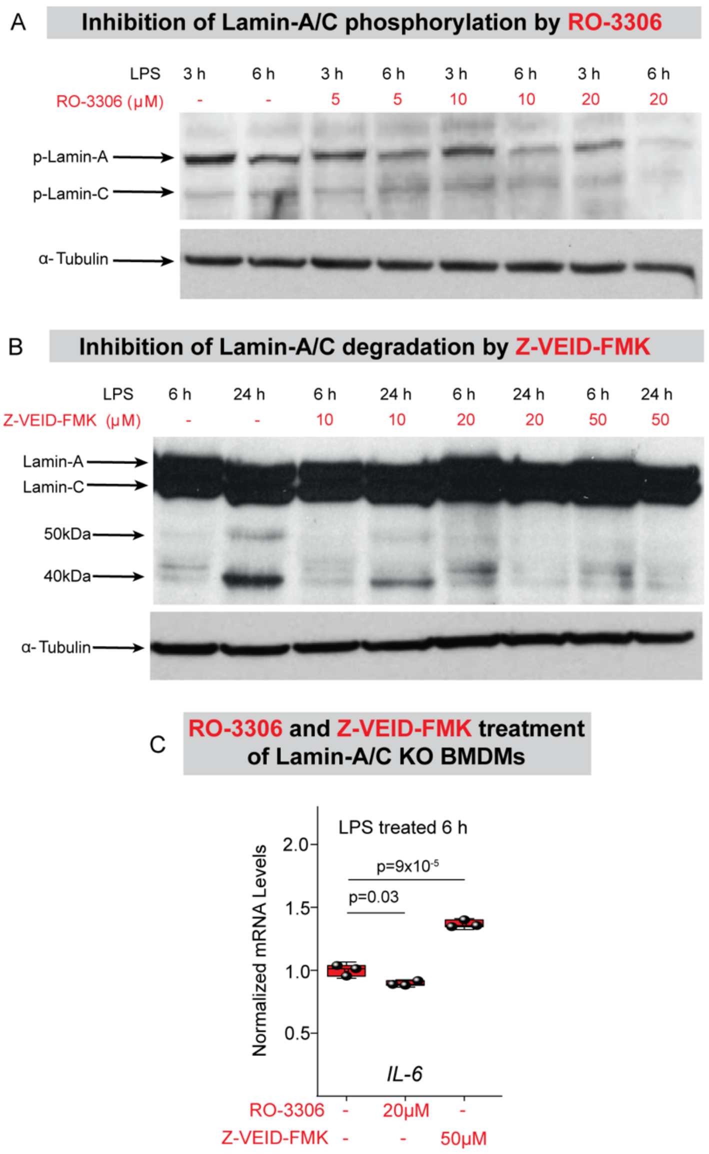


**Fig. S5: Lamin-A/C phosphorylation and degradation are inhibited upon RO-3306 and Z-VEID-FMK treatment, respectively: (A)** Immunoblot shows reduction in the total levels of p-Ser22-Lamin-A/C with RO-3306 treatment in wildtype BMDMs activated with LPS for 3 h and 6 h. α-tubulin served as a loading control. **(B)** Immunoblot shows inhibition of Lamin-A/C degradation with Z-VEID-FMK treatment in wildtype BMDMs activated with LPS for 6 h and 24 h. α-tubulin served as a loading control. **(C)** Box plots show gene expression levels of *IL-6* in LPS treated *Lamin-A/C* knockout (KO)-BMDMs also treated with RO-3306 and Z-VEID-FMK. Levels were normalized to LPS treated KO-BMDMs condition to find the fold change. In all the plots, the boxes show 25th and 75th percentiles, the middle horizontal line shows the median, small open squares show the mean, and whiskers indicate S.D. $p$ values were obtained with the two-sided Student´s $t$-test. All the experiments were independently repeated three or more times.

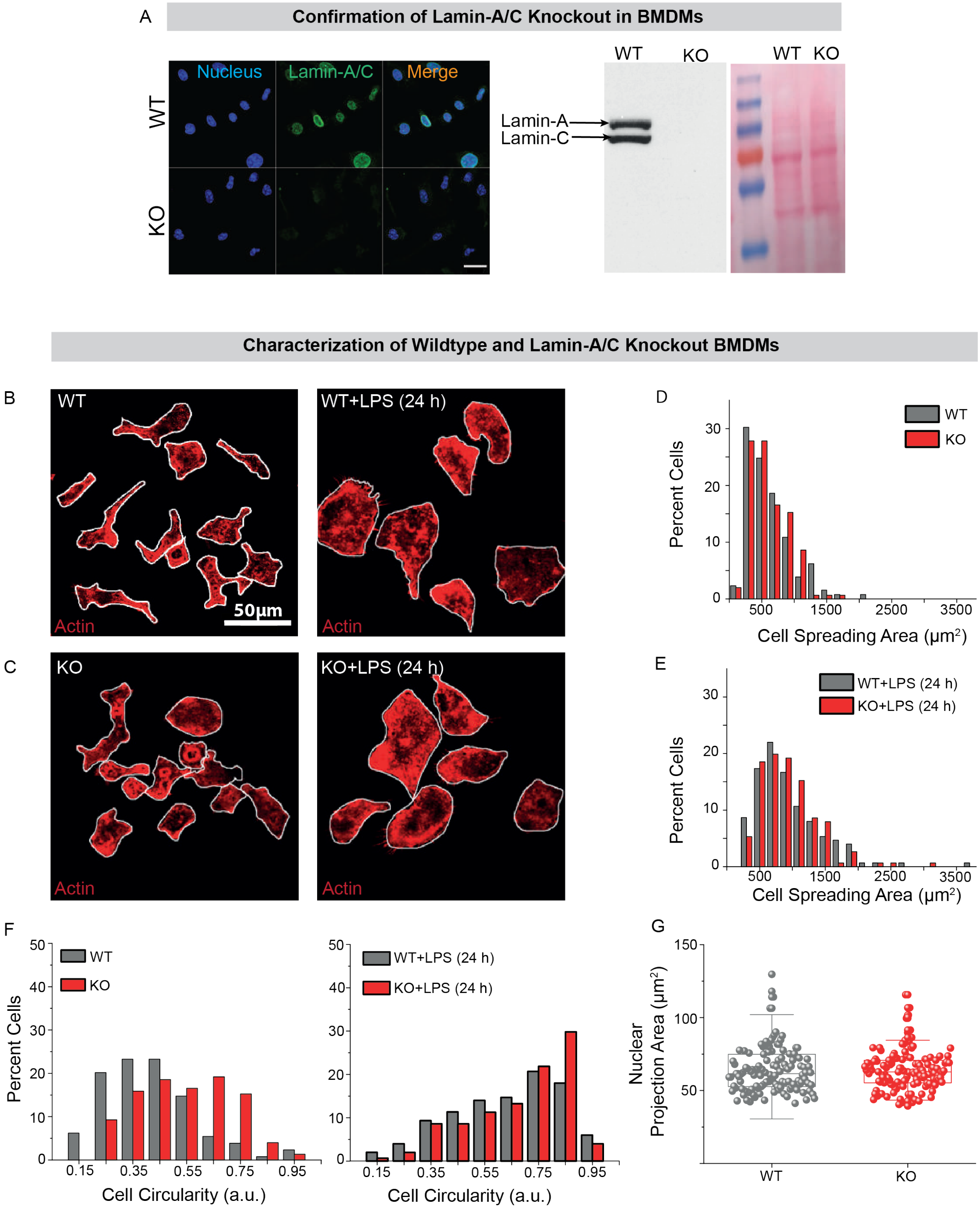


**Fig. S6: Characterization of Wildtype and *Lamin-A/C* knockout BMDMs: (A)** Representative image of Wild-type (WT) and *Lamin-A/C* knockout (KO) BMDMs stained for the nucleus (blue) and total Lamin-A/C (green), Scale bar = 20 μm **(Left)**. Immunoblot confirms absence of Lamin-A/C in KO-BMDMs. Also shown is the ponceau stain **(Right)**. **(B&C)** Representative images of WT- **(B)** and KO-BMDMs **(C)**, before **(Left)** and after LPS treatment for 24 h **(Right)**, stained for F-actin. Cell edges are marked in white. Scale bar = 50 μm. **(D&E)** Normalized distributions of cell spreading areas, before **(D)** and after LPS treatment for 24 h **(E)**, of WT- and KO-BMDMs. **(F)** Normalized distributions of cell circularity, before **(Left)** and after LPS treatment for 24 h **(Right)**. **(G)** Box plots show nuclear area in untreated WT- and KO-BMDMs. In all the box plots, boxes show 25th and 75th percentiles, the middle horizontal line shows the median, small open squares show the mean, and whiskers indicate S.D. All the experiments were independently repeated three or more times.

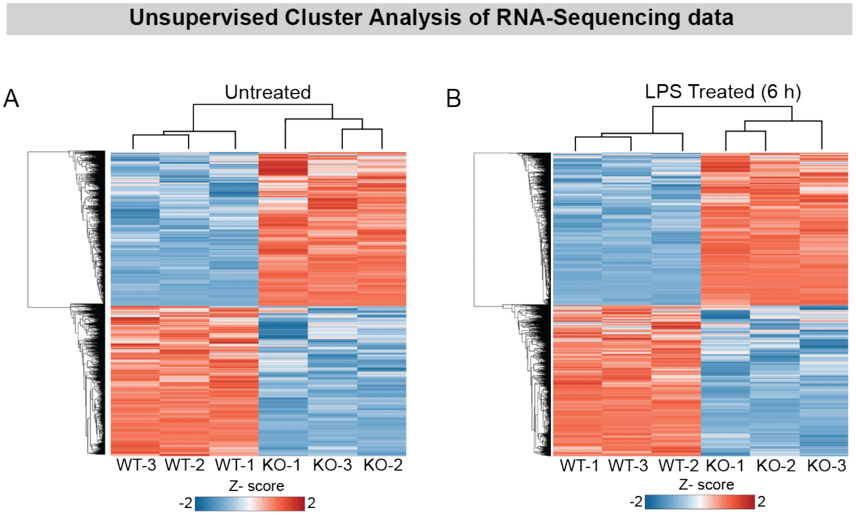


**Fig. S7: Unsupervised Cluster Analysis of RNA-Sequencing data: (A&B)** Heat maps cluster analysis of RNA-Seq data. Unsupervised cluster analysis on untreated **(A)**, and LPS treated **(B)** Wild-type (WT) and *Lamin-A/C*- knockout (KO)-BMDMs show separately clustering of WT- and KO-BMDMs. Each column represents one sample, each row a gene. Gene expression is represented in red for high expression, blue for low expression. Triplicate samples were used for RNA-Sequencing.

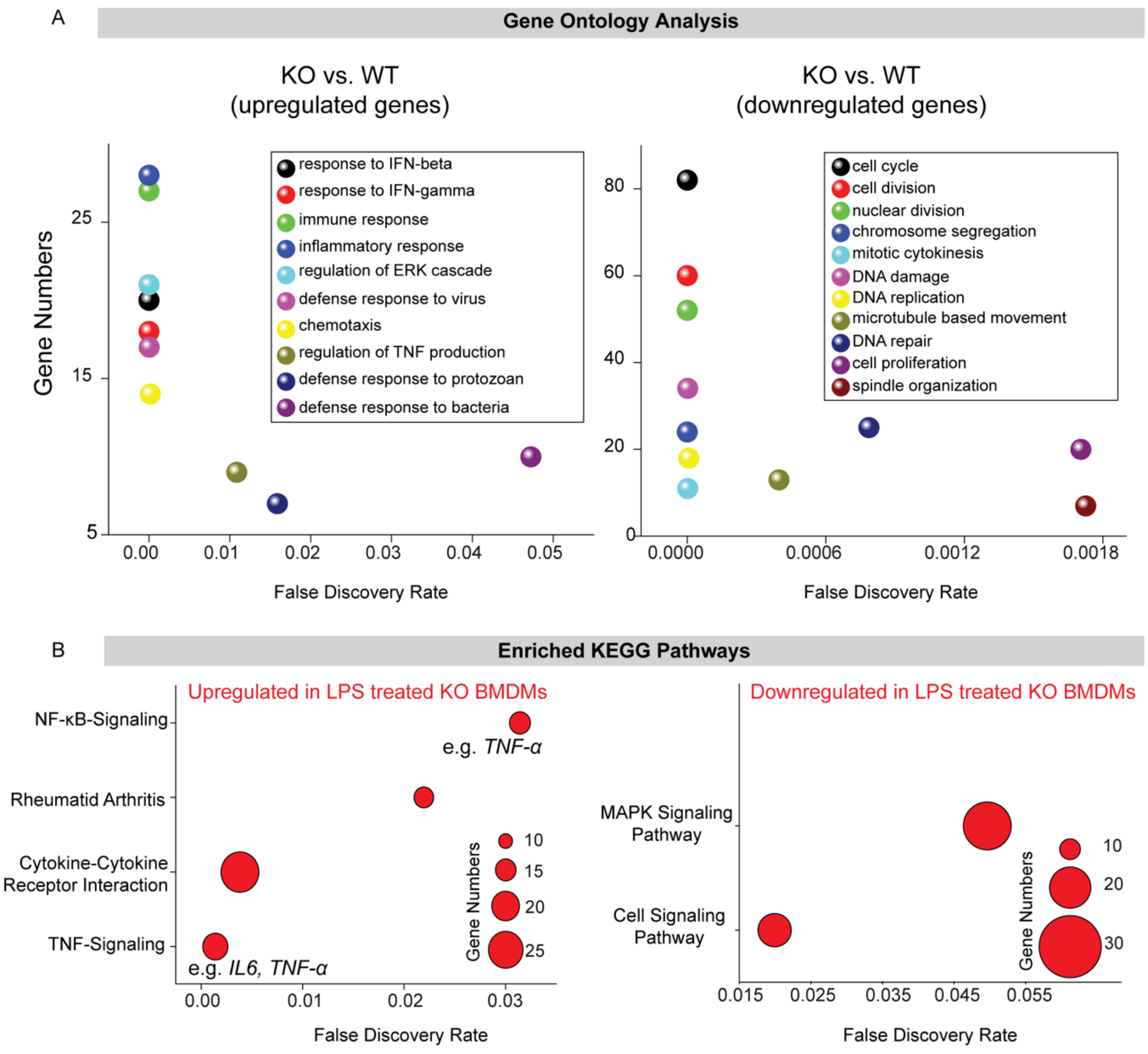


**Fig. S8: Gene Ontology analysis of upregulated and downregulated genes in *Lamin-A/C* knockout (KO) BMDMs and KEGG Pathways analysis of genes upregulated and downregulated in LPS treated KO-BMDMs: (A)** Gene-ontology analysis of upregulated and **(Left)** downregulated genes **(Right)** (Fold change>2 and p value < 0.05) in Lamin-A/C knockout (KO) BMDMs as compared to wildtype (WT) BMDMs. **(B)** KEGG Pathway analysis for all genes upregulated in LPS treated KO-BMDMs **(Left)** and all genes downregulated in LPS treated KO-BMDMs **(Right)** as compared to LPS treated WT-BMDMs.

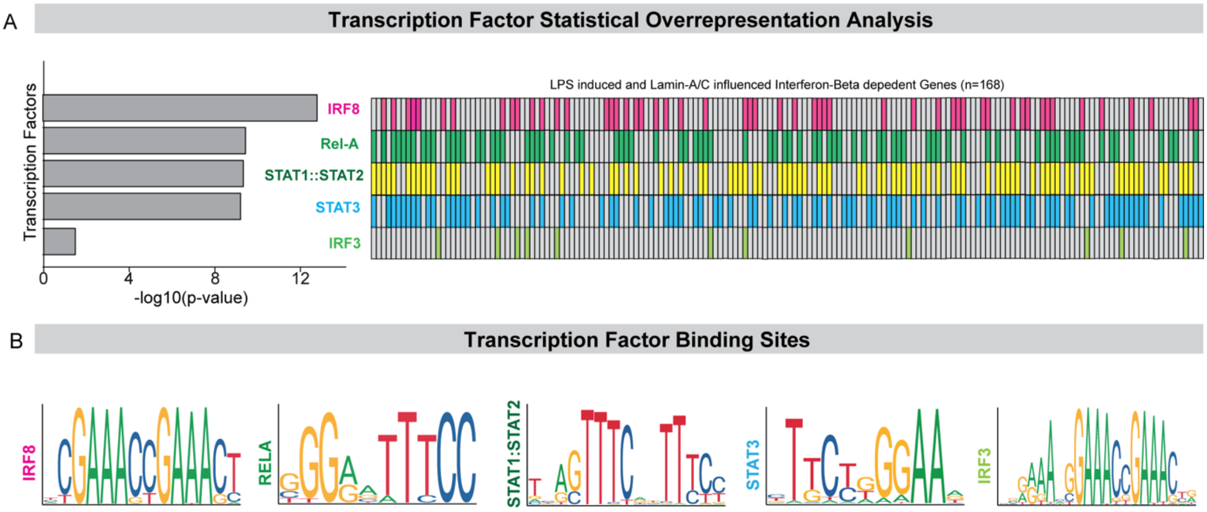


**Fig. S9: Transcription factor Statistical Overrepresentation analysis: (A)** Bar graphs show transcription factors with corresponding -log10(p-value) found by statistical overrepresentation analysis to detect enriched transcription factor binding sites in the promoter of the 168 genes (LPS induced, and Lamin-A/C regulated Interferon-β dependent Genes) **(Left)**. Heatmap shows the transcription factor binding for these 168 genes. Genes are arranged in descending order of their expression levels in LPS treated *Lamin-A/C*- knockout (KO) BMDMs **(Right)**. **(B)** Transcription factor binding sites for transcription factors found to be important for the transcription of the 168 genes (LPS induced, and Lamin-A/C regulated Interferon-β dependent Genes).

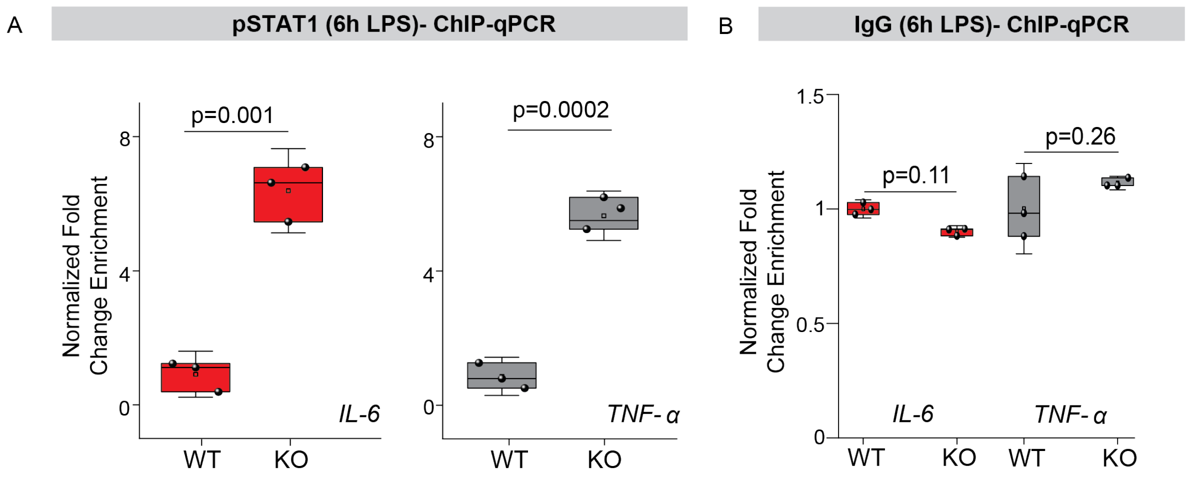


**Fig. S10: pSTAT1, and IgG ChIP-qPCR in LPS treated Wild type and *Lamin-A/C* Knockout BMDMs**: **(A)** Box plots show the ChIP-q-PCR analysis of pSTAT1 for the selected pro-inflammatory genes *IL-6* **(Left)** and *TNF-α* **(Right)** in LPS treated wildtype (WT) and Lamin-A/C knockout (KO) BMDMs. Levels were normalized to LPS treated WT-BMDMs condition. Data are plotted as normalized fold enrichment. Experiments were done in biological and technical replicates. **(B)** Box plots show the ChIP-qPCR analysis of IgG for the selected pro-inflammatory genes *IL-6,* and *TNF*-α for 6h LPS treated WT-BMDMs and KO-BMDMs. Levels were normalized to LPS treated WT-BMDMs condition. Data are plotted as normalized fold enrichment. Experiments were done in biological and technical replicates. In all the box plots, boxes show the 25th and 75th percentiles, the middle horizontal line shows the median, small open squares show the mean, and whiskers indicate S.D. For all the plots, *p* values were obtained with the two-sided Student´s *t*-test. All the experiments were independently repeated three or more times. ChIP-qPCR normalization has been explained in more detail in the Materials and Methods section.

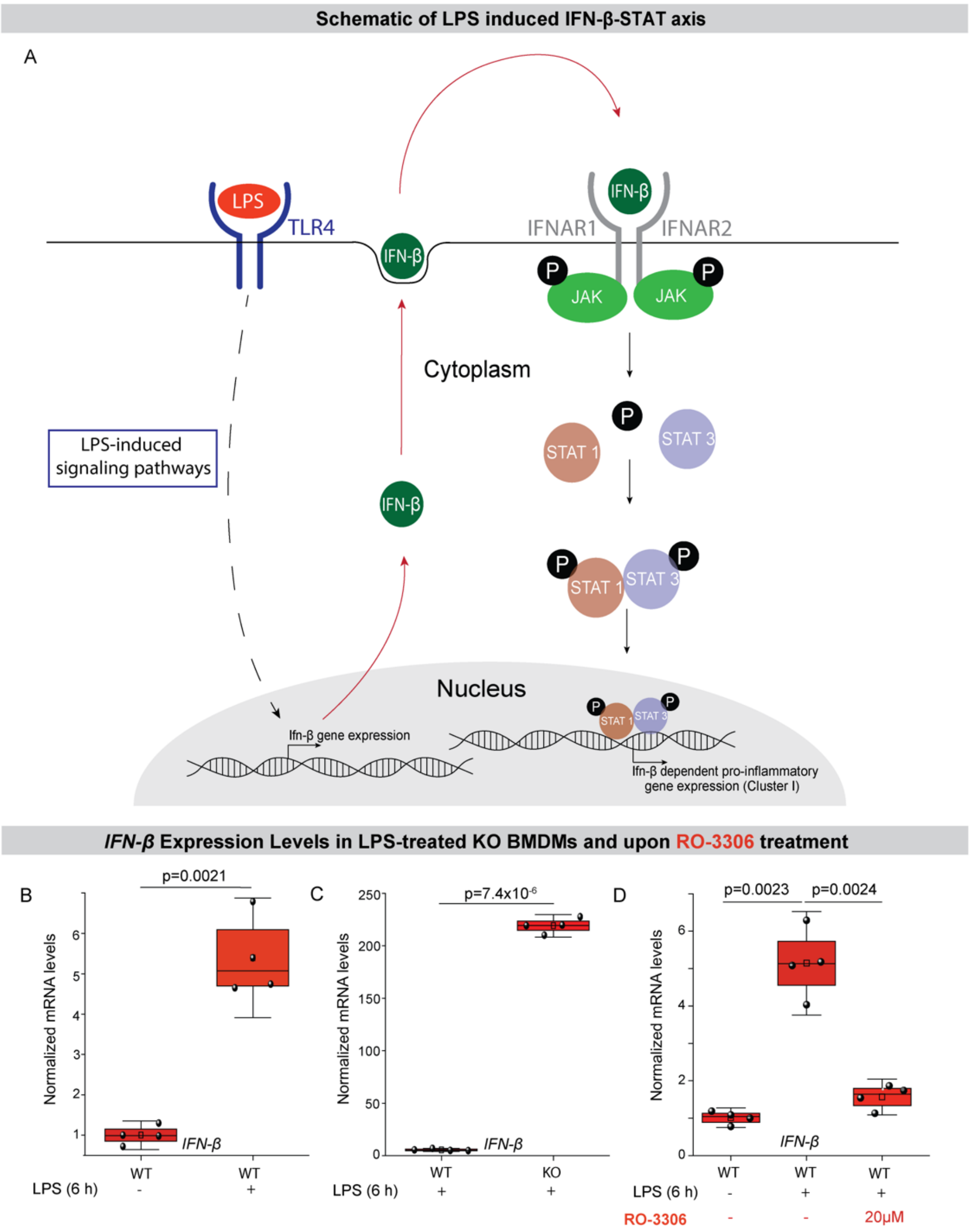


**Fig. S11: *IFN-β* expression levels in LPS-treated Wildtype, and *Lamin-A/C* KO-BMDMs, and upon RO-3306 treatment: (A)** Scheme of LPS-induced increased IFN-β secretion and subsequent activation of IFN-β-STAT axis resulting in increased pro-inflammatory gene expression. **(B)** Box plots show the differences in the mRNA expression levels of *IFN-β* in wildtype (WT) and WT+LPS treated BMDMs. Levels were normalized to untreated WT BMDMs condition to find the fold change. **(C)** Box plots show the differences in the mRNA expression levels of *IFN-β* in WT+LPS treated and *Lamin-A/C* knockout (KO)+LPS treated BMDMs. Levels were normalized to untreated WT BMDMs condition to find the fold change. **(D)** Box plots show the differences in the mRNA expression levels of *IFN-β* in WT, WT+LPS, and WT+LPS+RO-3306 treated BMDMs. Levels were normalized to untreated WT BMDMs condition to find the fold change. In all the box plots, boxes show the 25th and 75th percentiles, the middle horizontal line shows the median, small open squares show the mean, and whiskers indicate S.D. For all the plots $p$ values were obtained with the two-sided Student´s $t$-test. All the experiments were independently repeated three or more times.

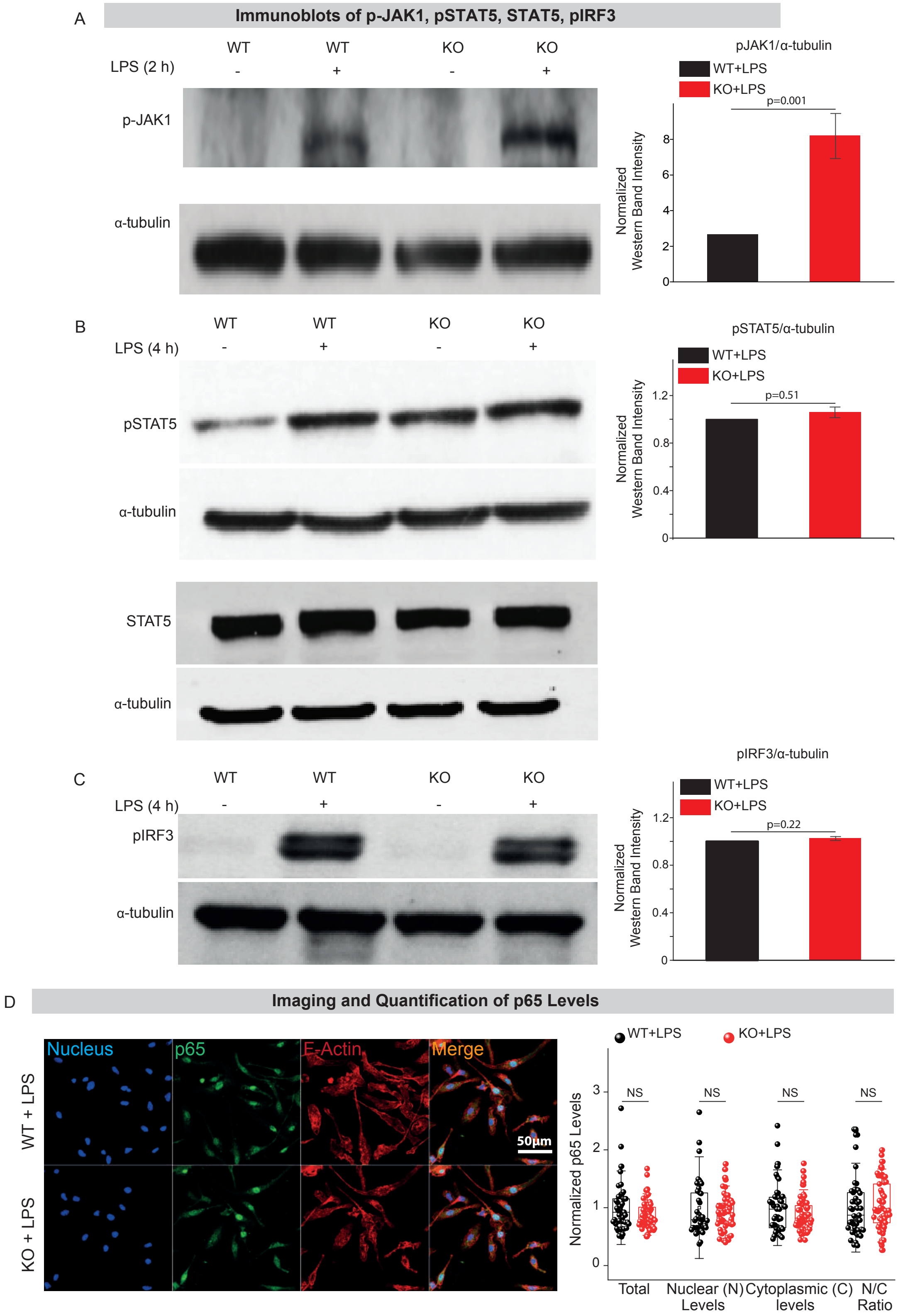


**Fig. S12: Quantifications of pJak1, pSTAT5, pIRF3 and p65 levels: (A)**Immunoblots show p-JAK1 levels in Wildtype (WT), WT+LPS, *Lamin-A/C* knockout (KO) and KO+LPS treated BMDMs. α-tubulin served as a loading control **(Left)**. Bar graph shows the normalized levels of pJAK1. pJAK1 levels were normalized to α-tubulin. Calculated values were normalized to the WT+LPS treated

BMDMs condition. Data are presented as Mean±S.E. **(Right)**. **(B)** Immunoblots show p-STAT5 levels (upper), and total-STAT5 (lower) levels in WT, WT+LPS, KO, and KO+LPS treated BMDMs. α-tubulin served as a loading control **(Left)**. Bar graph shows the normalized levels of pSTAT5. pSTAT5 levels were normalized to α-tubulin. Calculated values were normalized to the WT+LPS treated BMDMs condition. Data are presented as Mean±S.E. **(Right)**. **(C)** Immunoblots show levels of pIRF3 in WT-, WT+LPS-, KO- and KO+LPS-treated BMDMs. α-tubulin served as a loading control **(Left)**. Bar graph shows the normalized levels of pIRF3. pIRF3 levels were normalized to to α-tubulin. Calculated values were normalized to the WT+LPS treated BMDMs condition. Data are presented as Mean±S.E. **(Right)**. **(D)** Representative images of LPS treated WT- and KO- BMDMs stained for the nucleus (blue) total p65 (green), and F-actin (red). Scale bar = 50 μm **(Left)**. Box plots show the normalized levels of total p65, nuclear (N) p65, cytoplasmic (C) p65 and Nuclear to Cytoplasmic ratio (N/C) of p65 between LPS treated WT- and KO-BMDMs. Levels were normalized to LPS treated WT BMDMs condition to find the fold change. **(Right)**. In all the box plots, boxes show the 25th and 75th percentiles, the middle horizontal line shows the median, small open squares show the mean, and whiskers indicate S.D. For all the plots *p* values were obtained with the two-sided Student´s *t*-test. All the experiments were independently repeated three or more times.

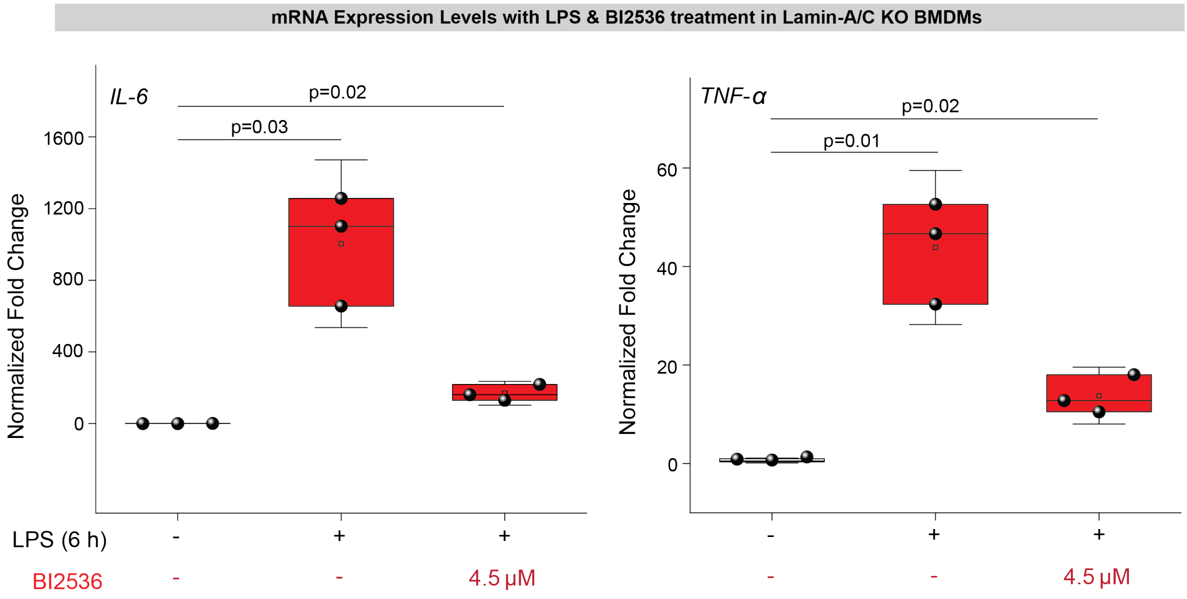


**Fig. S13: *IL-6* and *TNF-α* expression levels in LPS- and BI2536 treated *Lamin-A/C* Knockout BMDMs**: Box plots show the differences in the mRNA expression of *IL-6* **(Left)** and *TNF-α* **(Right)** in KO, KO+LPS, and KO+LPS+BI2536 treated BMDMs. Levels were normalized to untreated KO BMDMs condition to find the fold change. In all the box plots, boxes show the 25th and 75th percentiles, the middle horizontal line shows the median, small open squares show the mean, and whiskers indicate S.D. For all the plots $p$ values were obtained with the two-sided Student´s $t$-test. All the experiments were independently repeated three or more times.
